# Muscle growth by sarcomere divisions

**DOI:** 10.1101/2024.12.18.629106

**Authors:** Clement Rodier, Ian D. Estabrook, Eunice HoYee Chan, Gavin Rice, Vincent Loreau, Stefan Raunser, Dirk Görlich, Benjamin M. Friedrich, Frank Schnorrer

## Abstract

The sarcomere is the elementary contractile unit of muscles. Adult muscle cells chain thousands of sarcomeres into long periodic myofibrils that attach to the skeleton. How new sarcomeres are added during muscle growth is unknown. By live imaging and high-throughput image analysis, we have now tracked sarcomeric components during *Drosophila* muscle development and discovered that individual sarcomeres divide along the myofibril tension axis into daughter sarcomeres. This way, new sarcomeres can be inserted into contractile and mechanically intact myofibrils. We propose that sarcomere division is triggered by tension and local sarcomere damage originating from skeletal growth and muscle contractions. Sarcomere divisions repair damaged sarcomeres, ensure their mechanical integrity and synchronise sarcomere addition with skeletal growth during animal development.

## Main Text

Sarcomeres, the contractile units of muscles, have a stereotypic length of 2-3 micrometres in mammals (*1*). Muscle cells can be centimetres long, with thousands of sarcomeres connected in series to form myofibrils. These bridge across the entire cell, and their contraction moves the skeleton (*2*). How new sarcomeres are added into existing myofibrils, which are continuously under tension during development (*3*), is unknown. During early mammalian development, the fetal cardiomyocytes and skeletal muscle cells assemble sarcomeres to pump the fetal blood and to move its skeleton. These fetal cells are generally small, scaling the small size of the fetal heart and skeleton. As development proceeds, the distance between skeletal elements as well as the heart size enlarges, which is accompanied by a matched length increase of the skeletal muscle cells (muscle fibres) or, postnatally, of the cardiomyocytes (*2*, *4*, *5*). Since sarcomere length, which is measured between the two bordering Z-discs (Fig. S1A), is stereotypic and remains constant during this growth, a large number of additional sarcomeres need to be added to each existing myofibril. This addition represents a nontrivial problem as myofibrils are continuously under large mechanical tension and thus a myofibril would snap if opened in an uncontrolled way (*3*, *6*).

To date, the mechanism for sarcomere addition is unknown. One prominent hypothesis, the “end-hypothesis”, proposed that new sarcomeres are added exclusively at the terminal ends of myofibrils, where these are stably attached to tendon cells (*7*, *8*). One reason that made this hypothesis attractive is that the terminal Z-disc of each myofibril is thicker and, in some cases, has a zig-zag shape (*9*, *10*), which may allow the insertion of material without breaking the myofibrillar chain (*11*, *12*). However, to our knowledge, there is no direct observational evidence supporting this model. Visualizing sarcomere addition in growing muscles would be conclusive evidence, but this is hard to obtain in a developing mouse or human being. To this end, we reasoned that the problem is more fundamental and more ancient than mammalian evolution, as the basic sarcomeric architecture is highly conserved (*13*), and turned to *Drosophila* for studying the process.

### Flight muscles add sarcomeres rapidly

*Drosophila* dorsal-longitudinal indirect flight muscles (here called flight muscles for short) are indeed an ideal model as they are composed of only 6 large muscle cells in each hemi-thorax. After stably attaching to tendons at about 32 h after puparium formation (APF), each flight muscle cell assembles all of its 2000 myofibrils at once (*14*, *15*). Then, the muscle cells grow and double their length within a few hours of development (Fig. 1A)(*14*, *16*, *17*). To study the process of sarcomere addition in detail, we first quantified the length of the flight muscle fibres and the respective number of sarcomeres every 2 hours of development (Fig. 1B-D). We verified that individual sarcomere length remains constant at about 2 µm, while sarcomere regularity increases (Fig. 1C) as had been described (*18*). This analysis identified a particularly fast sarcomere addition phase between 34 h and 38 h APF, with one new sarcomere being added to each myofibril every two minutes (Fig. 1D).

**Fig. 1.**
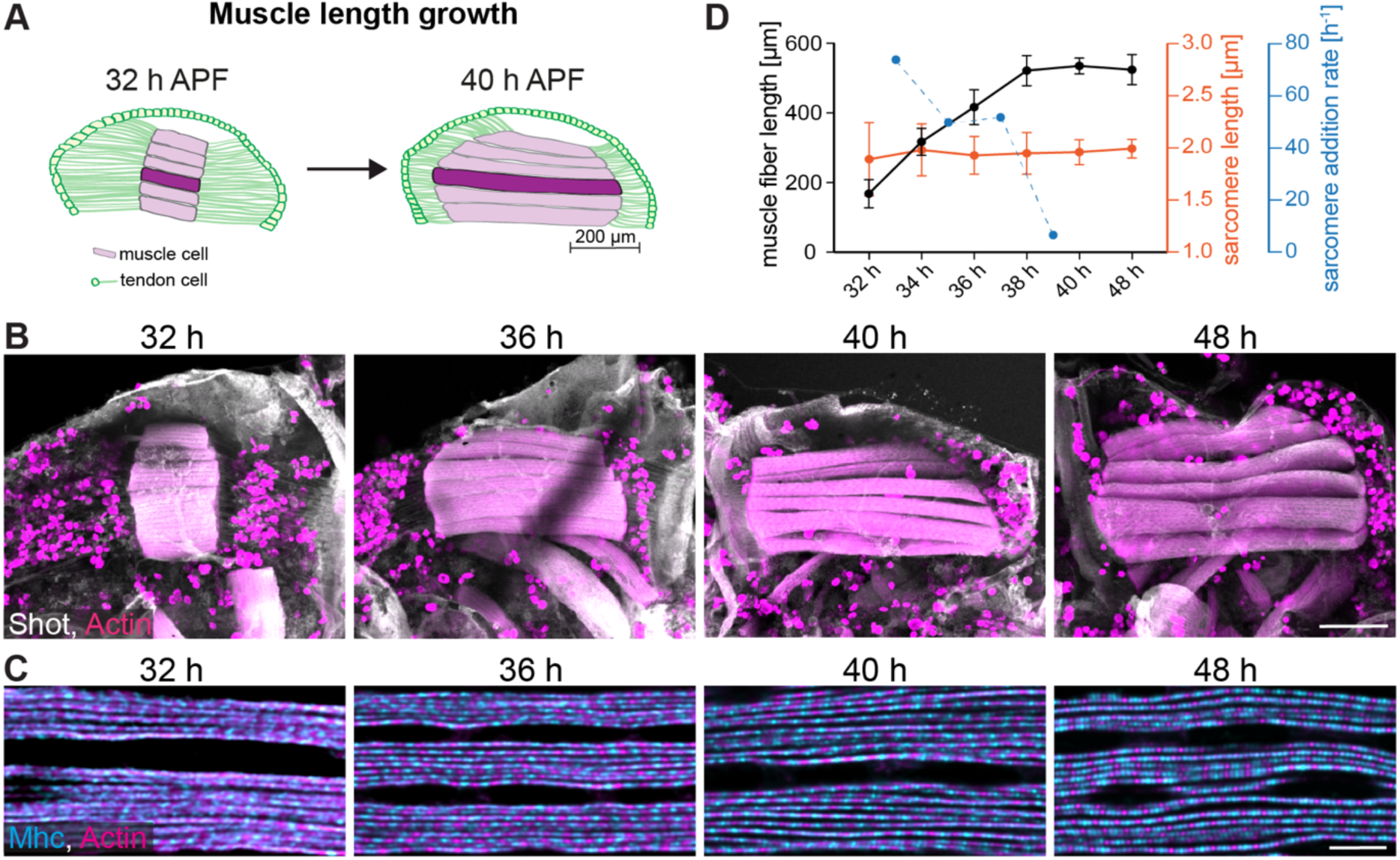
Muscle growth and sarcomere addition. **(A)** Schematic of a *Drosophila* hemithorax at pupal stages with the 6 dorsal-longitudinal flight muscles (DLMs) in magenta and the tendon cell epithelium in green. DLM4 is highlighted and was used for quantifications in (D). Note the stable connection of the flight muscles to the tendon cell extensions (green lines) during the growth phase from 32 h to 40 h APF. (**B, C)** Confocal images of pupae of the indicated stages displaying the six flight muscles stained for actin (phalloidin in red) and Shot (anti-Shortstop in grey) in (B). High magnifications displaying the myofibrils stained for actin (red) and myosin (anti-Mhc in blue) in (C). Scale bars are 100 µm in (B) and 5 µm in (C). Note the dramatic muscle length increase after 32 h APF. **(D)** DLM4 fibre length plotted in black and sarcomere length in red. Calculated sarcomere addition rate per myofibril per hour in blue. Error bars indicate standard deviation (s.d.). Number of scored sarcomeres is > 500 from 4 to 15 animals per stage (see Data S1).

If the end hypothesis of sarcomere addition would apply in this system, one would expect to find highly irregular newly added sarcomeres close to the terminal Z-discs. To test this, we performed high-resolution imaging, visualizing actin, myosin and titin (called Sallimus, Sls, the I-band titin in *Drosophila* (*19*)) at terminal and central muscle regions at 32 h, 36 h and 40 h APF (Fig. S1A-C). We indeed found thicker terminal Z-discs, judging from the actin and titin intensities; however, no obvious differences in sarcomere length or regularity were present compared to central muscle regions (Fig. 1B, C). To investigate the dynamics of the terminal sarcomeres directly, we imaged the growing muscle ends in living pupae using a titin-GFP (Sls-GFP) strain. We found that the terminal Z-discs can be traced over time without any obvious dynamics that may indicate sarcomere addition. Furthermore, no flow of sarcomeres away from the end can be observed in kymographs (Fig. S2A, B, Movie S1). The latter would be expected if new sarcomeres are only inserted at the terminal ends of the myofibrils. We found the same when imaging a myosin heavy chain-GFP (Mhc-GFP) strain tracing the terminal myosin filament stacks (Fig. S2C, D, Movie S2). Hence, we conclude that flight muscles rapidly add new sarcomeres during their fast growth phase, but identifying the position of addition prompted a more detailed investigation.

### Sarcomeres divide by segregating their myosin filament stacks

To identify possible sites of sarcomere addition, we developed an automated tool to trace myofibrils in 3D confocal stacks of flight muscles at pupal stages (Fig. 2A, see Methods, Fig. S3, Movies S3, 4). It automatically quantifies sarcomere length as well as protein distributions for the key sarcomeric components actin (labelling the actin filaments), titin N-term (Sallimus N-term, labelling the Z-discs), myosin (labelling the myosin filaments) and Obscurin (marking the M-band at the center of the myosin filaments, see Fig. S1A) in an unbiased way at 36 h and 40 h APF (Fig. 2B, C). We found the expected increase in peak regularity from 36 h to 40 h APF and the expected 2 µm sarcomere length, when measuring the distance between neighbouring Sallimus peaks representing the Z-discs (Fig. 2C). This showed that the 3D-detection method is accurate.

**Fig. 2.**
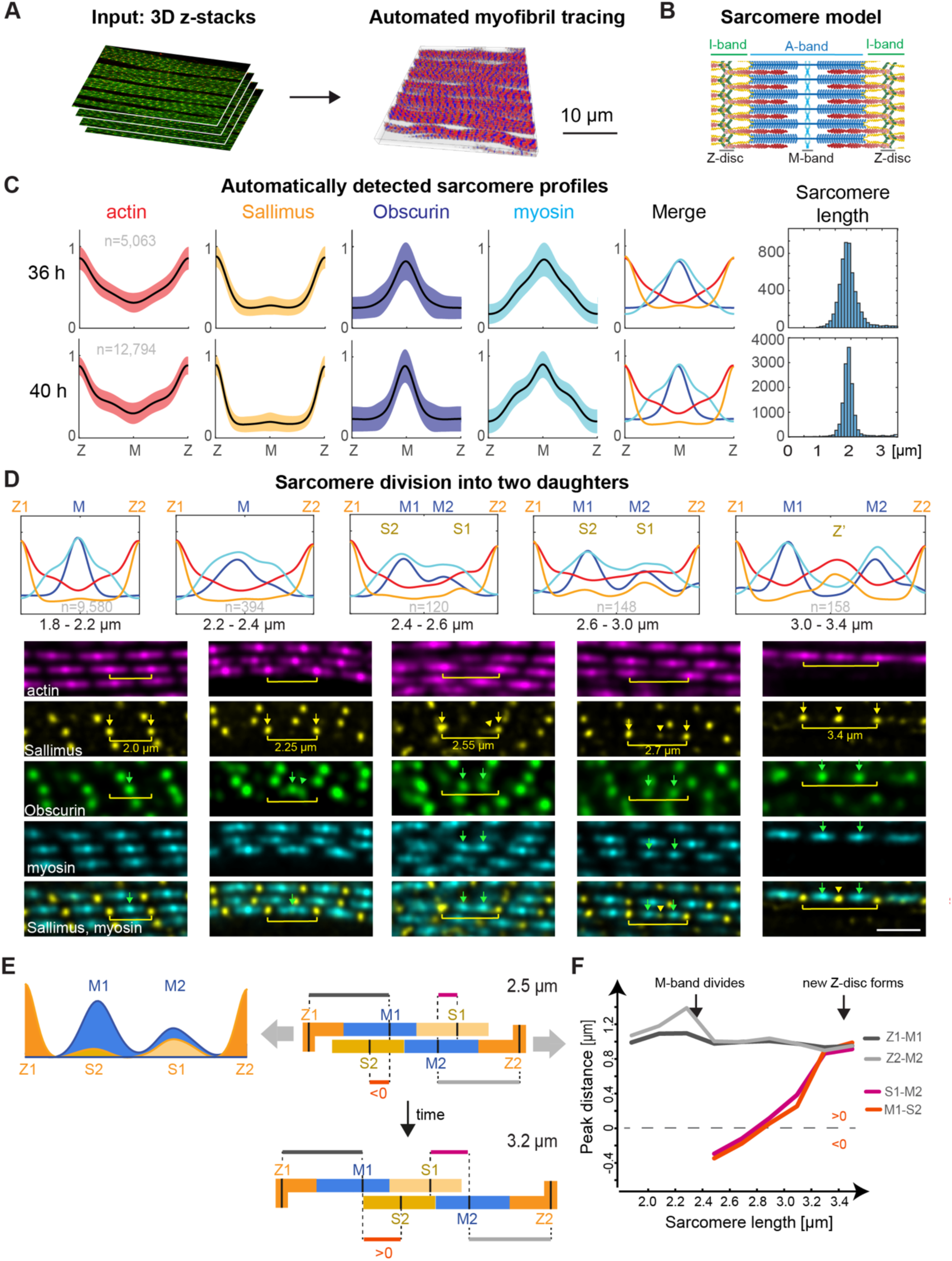
Sarcomere detection reveals sarcomere division by myosin segregation. **(A)** Summary of automated myofibril tracing from a confocal z-stack of flight muscles stained for sarcomere proteins (see Methods, Fig. S3, Movies S3 and 4). (**B)** Sarcomere scheme with the key regions labelled, see Fig. S1A for details). **(C)** Normalised intensity profiles computed from detected sarcomeres of actin (phalloidin in red), Sallimus (Sls-Nano2 in orange), Obscurin (anti-Obs in dark blue) and myosin (Mhc-GFP in light blue) shown as mean (black) ± s.d. (shaded regions) at 36 h and 40 h APF. 36 h APF: 5 samples, *n* = 5,063 sarcomeres; 40 h APF: 8 samples, *n* = 12,794 sarcomeres). Histograms of sarcomere length bins are shown to the right. **(D)** Top: Merged intensity mean profiles for the same proteins as in (C) sorted by sarcomere length bins using the 40 h APF flight muscles (*n* = 120 – 9,580 sarcomeres per bin, mean profiles from oriented individual profiles with the larger of the two Obscurin signals oriented to the left). Below are representative confocal images with highlighted sarcomeres matching the length categories. Note that two distinct myosin and Obscurin signals are visible in sarcomeres longer than 2.4 µm (green arrow and arrowheads); concomitantly Sls is moving away from the Z-discs (marked by yellow arrows) to form a new Z-disc (yellow arrowheads). Scale bar is 2 µm. **(E, F)** Peak-to-peak distances of Z1, Z2, S1, S2 (Sallimus) and M1, M2 (Obscurin) peaks computed from the mean intensity profiles shown in (D). Distances Z1-M1 and Z2-M2 (in grey) do not change when sarcomere length increases, while the distances M1-S2 and M2-S1 increase with slope ≈1 (F), suggesting that daughter sarcomeres slide relative to each other as rigid blocks (E, see also Movie S5).

Having validated the detection method, we next searched for deviations from the regular sarcomere pattern, with the motivation that such deviations may indicate sarcomere addition events. We scored sarcomeres with a length between 1.8 and 2.2 µm between neighbouring Z-discs as ‘regular’ sarcomeres. Additionally, we found many longer sarcomeres (8.6% of 10,479 sarcomeres), which display unusual, yet consistent protein distribution profiles (Fig. 2D, Fig. S4A): sarcomeres with a length between 2.2 - 2.4 µm show broader myosin and Obscurin peaks, while sarcomeres ranging from 2.4 - 2.6 µm in length display two distinct Obscurin and two distinct myosin peaks (labelled M1 and M2 in Fig. 2D). Seeing these data, we hypothesised that the fixed snapshots of increasing sarcomere length are reflecting a temporal event sequence. This would suggest that the widening myosin filament stack (called the A-band, see Fig. 1B) is segregating into two daughter myosin stacks along the tension axis, one to the anterior and one to the posterior side of the muscle fiber. Consistent with this hypothesis, we found that some Sallimus becomes detectable away from the Z-discs, as visible in the stainings as well as in the intensity profiles of the long sarcomeres (labelled as S1 and S2 in Fig. 2D, E). This suggests that some Sls was pulled out of the Z-discs towards the centre. When sarcomere length increases further (>2.6 µm), the myosin and Obscurin stacks move further apart and their intensity increases. At the same time, the central Sallimus populations assemble into a new Z-disc (called Z’) in the centre between the two daughter sarcomeres (Fig. 2D). Together, these snapshots are consistent with the hypothesis that a flight muscle sarcomere can divide into two daughter sarcomeres by segregating its myosin filament stacks along the myofibril tension axis (*15*).

As the suggested sarcomere divisions are likely induced by high tension created by growth of the pupal thorax, we wondered how the division is molecularly controlled to prevent a fatal rupture of the sarcomere. Sarcomeres are mechanically held together by titin springs linking the Z-disc to the myosin filaments. *Drosophila* contains two titin homologs, Sallimus and Projectin, with Sallimus linking the Z-disc to the end of the myosin filaments, whereas Projectin is located at the ends of the myosin filaments in flight muscles (see Fig. S1A)(*19*). Thus, segregating the myosin filaments into two distinct blocks should either break the Sallimus/myosin linkage or the Sallimus/Z-disc linkage. Following the pseudo-temporal sequence of our detected intensity profiles and quantifying protein amounts in a given sarcomere length bin (Fig. S5) argues for a combination of both. Thus, localised sarcomere damage triggers the sarcomere dvisions. In most cases, the mother sarcomere divides asymmetrically with a larger myosin block (labelled M1) moving to one side, here left, together with a pronounced Sallimus signal originating from the right Z-disc (S1), while the remaining Sallimus stays at the Z-disc (Z2). At the same time, a smaller myosin block (M2), moving right, segregates together with a smaller Sallimus signal (S2) from the left Z-disc (Z1) (Fig. 2D - F, Fig. S5). As sarcomere length continues to increase, the mutual distances between Z1, M1 and S1 remain constant, suggesting that these move as an intact block. The same is the case for a second block consisting of Z2, M2, S2. The distance between M1 and S2 (and thus between the two blocks) continuously increases, exactly as sarcomere length increases (Fig. 2E, F, Movie S5). Additionally, we find that Projectin segregates together with the myosin filaments (Fig. S6). During a division, the left and right neighbouring sarcomeres remain intact, apart from the division-induced reduction of actin and Sls intensities at the shared Z-discs (Fig. S4B). From these quantitative data, we conclude that defined local damage of protein interactions allows the two daughter sarcomeres to segregate as two blocks, likely sliding past each other, until the two daughter Z-discs S1 and S2 are brought into contact and subsequently merge to build a new Z-disc separating the two daughter sarcomeres. The further the division process advances, the more newly synthesised protein components are recruited to replenish the available binding sites on the daughter sarcomeres, which turn indistinguishable from their neighbours after reaching a mature sarcomere length of 2.0 µm. This mechanism is compatible with the observed relatively fast turnover rates of myosin, with 50% exchange in about 30 minutes in sarcomeres of 36 h APF pupae (Fig. S7, Movie S6).

### Sarcomere division – live

Thus far, we collected evidence for sarcomere divisions based on fixed images. To directly investigate the temporal dynamics of myosin filament stack segregation during sarcomere division in growing flight muscles, we performed *in vivo* live imaging using two-photon microscopy of an endogenously tagged functional Mhc-GFP line (*15*, *20*) (Fig. 3A). In these movies, we identified several events, in which a single myosin stack representing one A-band segregated into two separate stacks of lower intensities over the course of 10 to 20 minutes (Fig. 3A, B, Fig. S8A-C, Movies S7, 8). We quantified the distances between the adjacent myosin peaks along the myofibril to track the dividing myosin stack in comparison to its neighbours (Fig. 3B, C). As soon as the myosin stack divided, its two daughters were pulled away from each other at a speed of about ∼0.1µm / min until their distance had increased to about 1.8 µm, which is close to the mature sarcomere length (Fig. 3C). In the course of the division, the intensities of the daughter myosin peaks increased, again suggesting that new myosin proteins are recruited to replenish protein levels in the daughter sarcomeres (Fig. 3B). These data strongly support the sarcomere division mechanism that we identified based on the fixed images (Fig. 2).

**Fig. 3:**
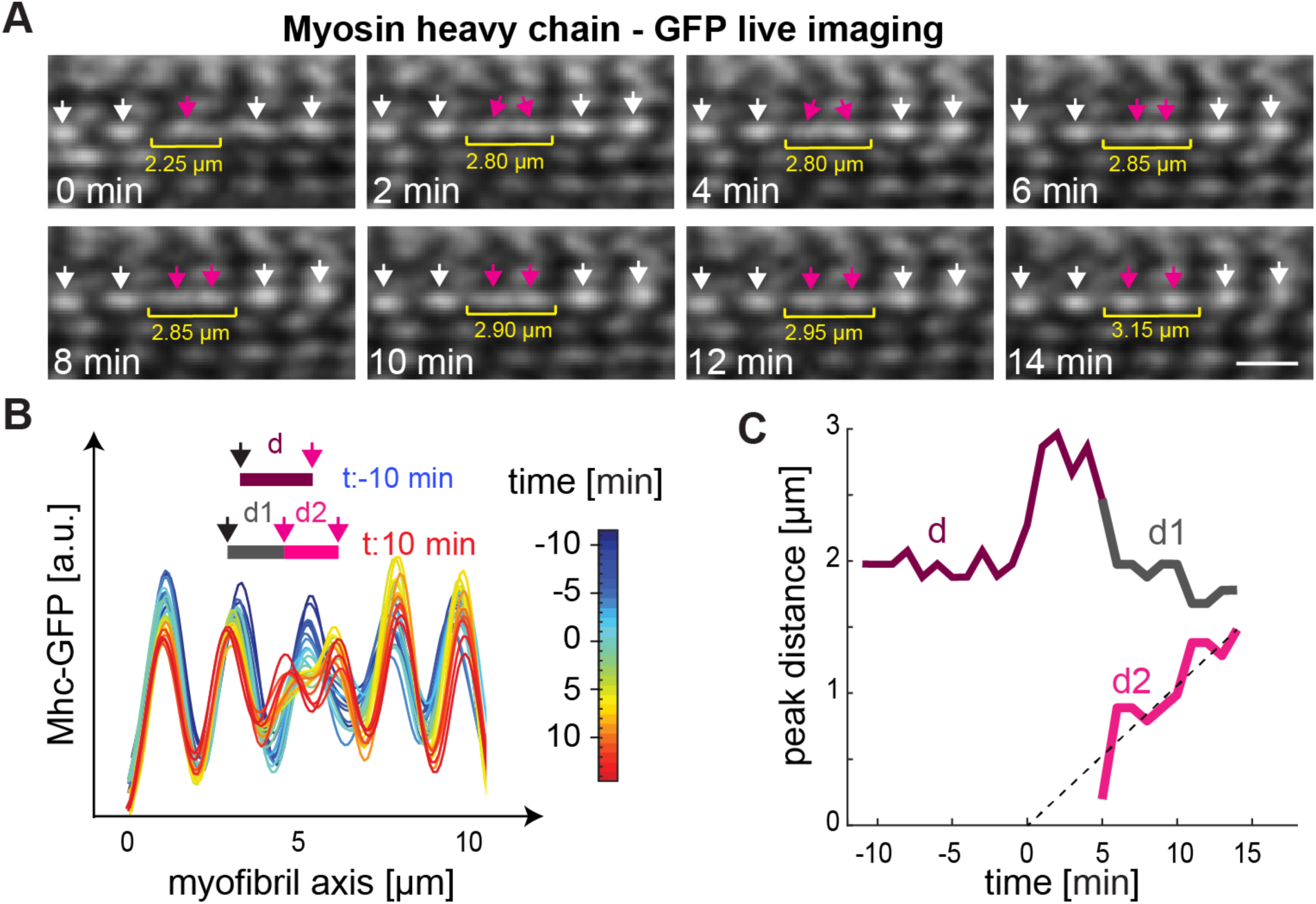
Live imaging of sarcomere division. **(A)** Stills from time-lapse movie of Mhc-GFP expressing flight muscles of a 36 h APF pupa acquired with two-photon microscopy (see Movie S7). Arrows indicate the myosin filaments of one selected myofibril. Sarcomere length is indicated with the yellow brackets. The dividing myosin filament stack is labelled with magenta arrows. Note that the increase in sarcomere length correlates with the segregation of the two new myosin peaks. Scale bar is 2 µm. (**B)** Mhc-GFP intensity profiles along the highlighted myofibril in (A) for different colour-coded time points. Distances between Mhc-GFP peaks are highlighted for two frames (t: −10 min in blue, t: 10 min in red). Time 0 minutes indicates the putative start of the division event. (**C)** Inter-peak distances of intensity profiles from panel B as function of time. Colour-code corresponds to inset in (B).

### Sarcomere division in larval muscles

Flight muscle sarcomeres are not cross-striated: their myofibrils are isolated (*21*) and hence sarcomeres in neighbouring myofibrils can divide independently. Furthermore, flight muscle myofibrils are not yet fully contractile during the pupal stages that we analysed above. To investigate if a similar sarcomere division mechanism also applies during growth of actively contractile cross-striated muscle types, we chose to investigate growth of the cross-striated *Drosophila* larval body muscles that resemble mammalian skeletal muscles more closely. At the end of embryogenesis, larval muscle sarcomeres are fully contractile and power the larval movements. During the three larval instars, the larvae significantly increase in length; the segment spanning ventral-longitudinal muscles increase from about 10 sarcomeres in small L1 larvae to about 80 sarcomeres four days later in fully grown L3 larvae (*22*, *23*), suggesting an addition rate of one new sarcomere every 1 - 2 hours. In contrast to immobile pupae, larvae move vigorously while eating food to fuel larval growth.

To document the larval sarcomere morphology in detail, we first applied plasma-focused ion beam scanning electron microscopy (pFIB/SEM) and imaged large volumes of larval muscle resulting in complete three-dimensional (3D) reconstructions. The SEM volumes revealed thin myofibrils that were stacked laterally, as expected (Fig. S9A). Interestingly, we also find that Z-discs and myosin filament stacks are frequently out of register laterally, with the Z-discs showing frequent Y-shaped branches (Fig. S9A, B). The high contrast of the Z-discs enabled us to generate detailed 3D reconstructions (Fig. S10A-C). We labelled the Z-discs at the edge of the myofibril stacks and segmented their densities towards the centre. This allowed us to trace the Z-discs, revealing an extensive, interconnected Z-disc network (Fig. 4A, Movie S9). These results inspired us to hypothesise that tensile forces may induce a progression of the Y-shape branches, similar to the opening of a zipper, which would be consistent with a tension-induced sarcomere division mechanism.

**Fig. 4:**
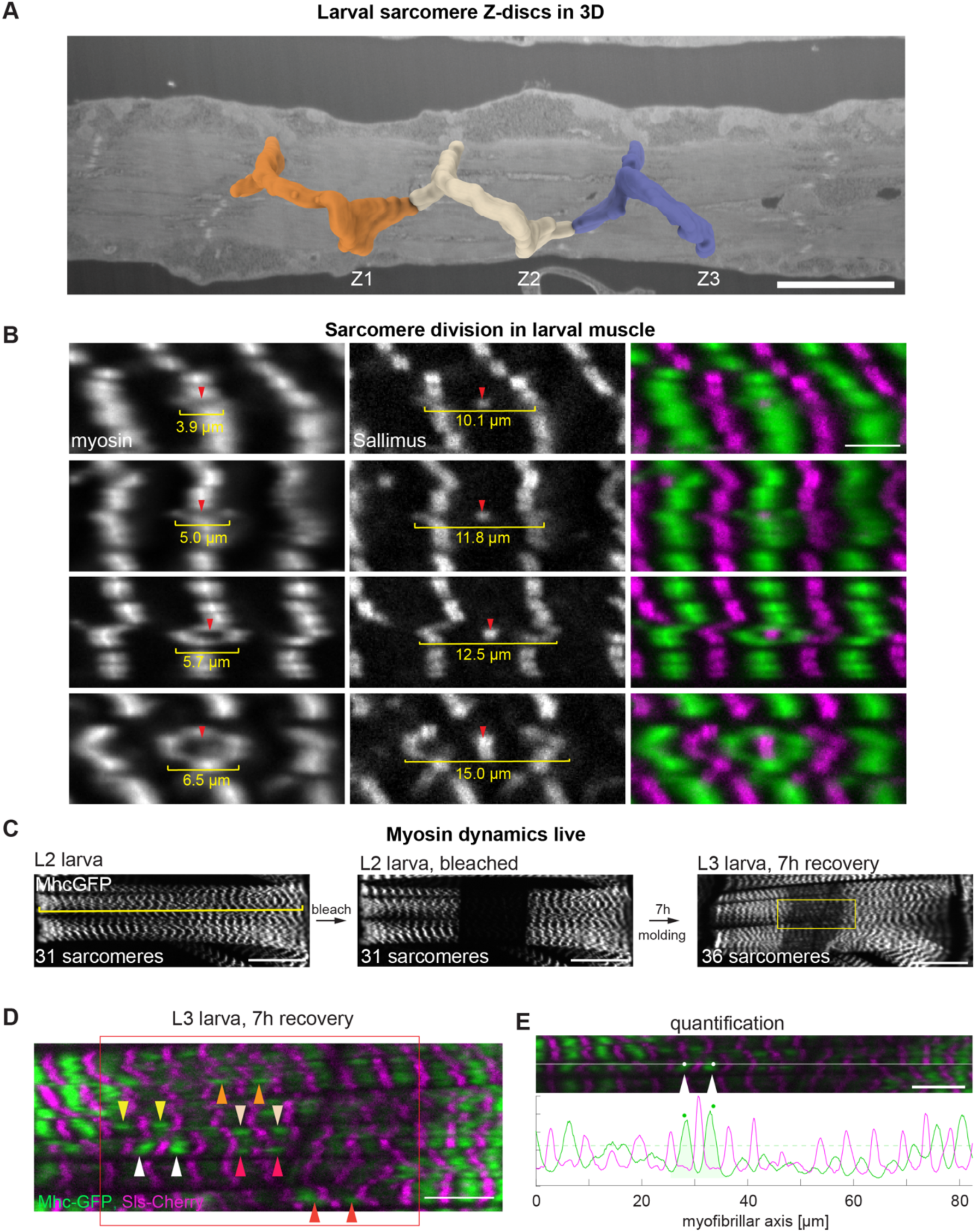
Dividing sarcomeres in larval muscles. **(A)** Scanning electron microscopy segmentation of larval Z-discs in a 3D volume starting from the muscle surface using the slices shown in Fig. S10. Each colour indicates voxels that were associated to one complete Z-disc at the lateral edge of the myofibril. See also Movie S9. Scale bar is 7 µm. **(B)** Living *Drosophila* larval muscles labelled with Mhc-GFP (green) and Sls-Cherry (magenta) display myosin ‘rings’ sorted according to increasing diameters indicated in yellow with increasing amounts of Sls in their centre marked by red arrowheads. Scale bar is 5 µm. **(C)** Mhc-GFP in living L3 larval muscle (dorsal oblique 2, DO2), before FRAP (left), after FRAP (middle, FRAP region marked by a red rectangle), as well as after 7 h recovery with feeding (right, approximate region shown as high magnification in (D) is marked with a yellow rectangle). Note that yellow line marks 31 sarcomeres before FRAP, which increases to 36 sarcomeres after 7 h recovery. Scale bar is 50 µm. **(D)** High magnification of yellow region in (C) with Mhc-GFP in green and Sls-Cherry in magenta. Note that Mhc-GFP does recover in pairs marked by arrowheads of the same colour (see also Movie S12). Scale bar is 10 µm. **(E)** Automatic quantification of recovered myosin peaks in the bleached region using parallel line scans along the myofibrillar direction (*upper*, white line shows example line scan), and corresponding intensity profiles of Mhc (green) and Sls (magenta) with identified pairs of recovered myosin peaks (*lower*, green dots).

To investigate such a possible mechanism in larval muscle in more detail, we stained larvae expressing Mhc-GFP with phalloidin to label the actin filaments and an N-terminal anti-Sls nanobody to label the Z-discs and imaged them with confocal microscopy followed by a deconvolution algorithm for optimal z-resolution. Generally, these markers reproduced the cross-striated larval sarcomere morphology. Consistent with our EM data, we frequently identified Z-discs as well as the myosin filaments as branched Y-shaped structures, sometimes even as double Y-shaped filament stacks that appear as “rings” in one plane (Fig. S11A, Movie S10).

To rule out that these structures could a be fixation artefact, we next analysed living intact anaesthetised larvae expressing Mhc-GFP or Obscurin-GFP to label the M-band, together with Sallimus-Cherry marking the I-band close to the Z-disc. We found the same myosin stack ring structures as in the fixed images. Analogous to the cross-striated flight muscles, we hypothesised that rings with a small diameter might be Y-shaped zippers that have only recently opened, while large rings might be developmentally older. Accordingly, we ordered the rings according to their diameter and assayed how much Sallimus is present in the ring centre. If the diameter of the ring is small, only little Sallimus is present in its centre, while with increasing ring diameter, the amount of central Sls increases to reach the Sls concentration of a regular sarcomere (Fig. 4B, Fig. S11B-D). At the same time, the total amount of Obscurin-GFP present in the segregated myosin stacks doubles, thus replenishing the segregated myosin filament stacks to normal Obscurin levels (Fig. S11B-D). These static images suggest a related sarcomere division mechanism as found in the flight muscles: M-bands, or possibly also Z-discs, segregate into two blocks, which eventually mature into two daughter sarcomeres separated by a new Z-disc or M-band in the centre of the new sarcomere. The Y-shaped morphology of the junction suggests that the “zipper-like” mechanism can proceed along the short axis of the muscles (transverse to the tension axis) and hence numerous new sarcomeres can be added throughout the large muscle fiber.

We next attempted to visualise the zipper dynamics in larval muscles directly. We imaged anaesthetised living larvae expressing Obscurin-GFP and tracked the M-bands over time. However, within one hour no obvious zipper mechanism was detectable under these conditions (Fig. S12A, Movie S11). This lack of sarcomere addition is likely caused by the reduced sarcomere contractions and the absence of muscle fibre growth during anaesthesia, consistent with a key role of larval cuticle growth, to which the muscles are connected, and hence tension exerted on myofibrils, to break protein bonds in the sarcomere and advance sarcomere addition.

To allow for normal muscle contractions and larval growth, we next anaesthetised the larvae only briefly and then placed them back into food. To still be able to track sarcomere proteins, we photo-bleached parts of one muscle to distinguish old from newly synthesised or newly recruited Mhc-GFP (Fig. S12B). When photo-bleaching Mhc-GFP throughout an entire z-volume of the muscle, the bleached muscle area could be distinguished from neighbouring regions for up to 4 days, while the larvae were feeding. Strikingly, when imaging the bleached area again, we found sites of high Mhc-GFP fluorescence often close to Y-shaped Mhc branches or as localised pairs, suggesting that newly synthesised Mhc-GFP was incorporated locally in neighbouring myosin stacks of two daughter sarcomeres that segregated recently and replenished their myosin content (Fig. S12B). This is consistent with the hypothesis that the sarcomere division can propagate at the Y-shaped stacks using a zipper mechanism.

However, the Mhc-GFP recovery under these conditions took days and the larvae grew only little. Thus, we repeated the experiment with milder anaesthesia and milder bleaching at late L2 stage larvae, which we placed back into the food. Some of these larvae moulded to the L3 stage within a few hours. Under these conditions, we indeed found an addition of 5 new sarcomeres in 7 hours matching the above estimated wild-type growth rate (Fig. 4D). Zooming into the recovered area revealed several pairs of high Mhc-GFP signal through the recovering volume, which occur with high statistical significance (p < 10^-4^, Fig. 4E, Movie S12). Together, these data provide strong evidence that cross-striated larval sarcomeres divide by segregating myosin filament stacks into daughter stacks anywhere throughout the growing muscle fiber.

The here identified a mechanism allows to add new sarcomeres into existing myofibrils anywhere along the growing muscle fibre. This is a fundamentally different mechanism compared to the end-hypothesis, which proposed that new sarcomeres are exclusively inserted at the terminal Z-discs of muscle fiber ends (*7*, *8*, *11*, *12*). The latter may still apply under certain circumstances but would require the transport of all newly synthesised sarcomere components to the muscle fiber ends and the damage to be limited to the ends.

How are sarcomere additions regulated? Every sarcomere division requires the controlled breakage of protein-protein interaction bonds, a controlled sarcomere damage, to allow segregation of the sarcomere components, without an uncontrolled break of the entire myofibril. We propose that a controlled damage is triggered by the high mechanical tension, which is built-up by skeletal growth that stretches the attached muscles, while they keep contracting. As a consequence, tensile forces in the sarcomere increase, until a threshold is passed and a sarcomere division is triggered or the zipper progresses forward. Which protein may sense this high tension to initiate the division? The best candidate is the giant I-band titin Sallimus, which in analogy to the mammalian titin is responsible for most of the passive sarcomere tension and is stretched across the I-band region, when a sarcomere is stretched (*3*, *24–26*). We propose a model in which high mechanical forces cause the breakage of some of the Sallimus – Z-disc linkages, or possibly also some Sallimus – myosin linkages, on both Z-discs flanking the dividing sarcomere. This allows the segregation of some bipolar myosin filaments to the right and some to the left, thus adaptively reducing tension within the myofibril (Fig. 5, step 1 – Division initiation). This division initiation creates free binding sites that allow the recruitment of new sarcomere proteins, making the division process irreversible (Fig. 5, step 2 – Novel protein recruitment). During the entire process, myosin can still interact with actin filaments, which prevents the uncontrolled rupture of the entire myofibril. Finally, the centrally located Sallimus, actin and Z-disc components establish a new Z-disc between the segregated myosin stacks (Fig. 5, step 3 – Establishment of a new Z-disc).

**Fig. 5:**
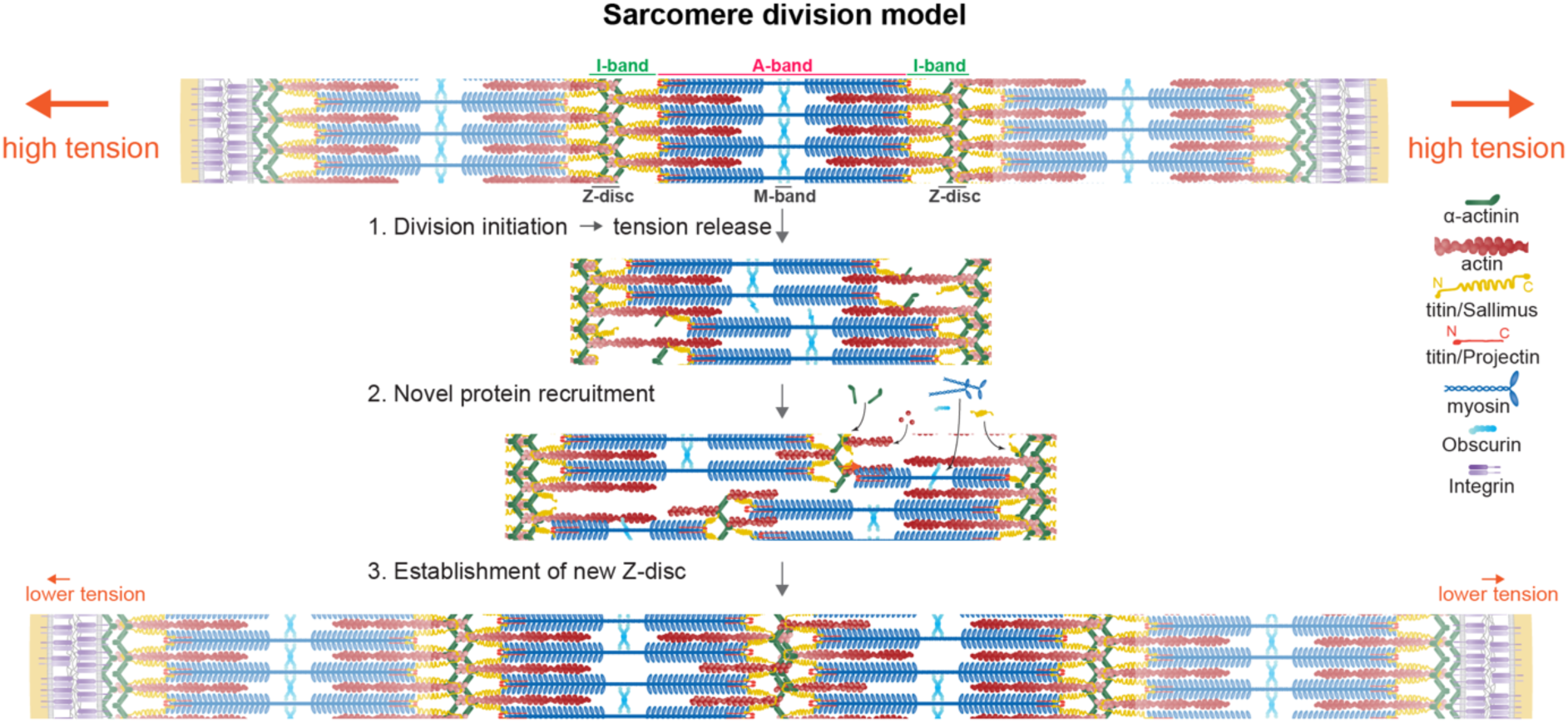
Sarcomere division model.

Working model of the tension-induced sarcomere division in *Drosophila* flight muscle sarcomeres. The growth of the skeleton, marked by integrin-based attachments, induces high tension, which triggers the division of the middle sarcomere. Thus, the sarcomere number increases from three to four. Details see text.

Is a comparable sarcomere division mechanism present in other striated muscles? Electron microscopy studies of growing crab muscles (*Callinectes sapidus*, an arthropod) showed structures that can be interpreted as dividing A-bands, consistent with our model (*27*). Hence, the here proposed sarcomere division mechanism is likely conserved in arthropods. In mammalian muscles, the mechanism may vary as sarcomeres should break at their weakest point. Mammalian sarcomeres contain in addition to Obscurin the strong M-band crosslinker myomesin, a protein not present in insects (*28*, *29*) and mammalian titin is tightly bound to myosin along the entire thick filament (*30*). If the myomesin crosslinks at the M-band as well as the titin – myosin interactions are more stable than the α-actinin crosslinks at the Z-disc, sarcomere division in mammalian muscle is likely initiated by a tension-induced separation of Z-discs. Such Z-disc “streaming” was frequently observed after either eccentric exercise or mechanical stretching of cardiomyocytes in culture; both protocols generated very high tension and thus local damage on sarcomeres (*31–34*). Since both tension protocols can result in the addition of sarcomeres within muscle fibers (*32*, *35*, *36*), a tension-induced sarcomere division mechanism may also occur during mammalian muscle growth.

## Acknowledgements

We thank Mathias Gautel and all his group members, as well as the Schnorrer, Raunser and Görlich groups for their stimulating discussions within the StuDySARCOMERE ERC synergy grant. We thank Olivier Pourquie and all members of the ‘Biological Algorithms’ Friedrich group for inspiring discussions within the HFSP network collaboration. We thank David Gross and Stefan Gumhold for their help with 3D-visualization. We thank Yasmin Magdy Emadeldin Mohamed Abdelghaffar for help with documenting the sarcomere detection tool. We are grateful to the IBDM fly and imaging facilities and especially to Pierre Mangeol for help with image acquisition, set-up, maintenance of the microscopes and helpful comments on this manuscript. The FIB-SEM sample preparation was performed on the PiCSL-FBI core facility (Nicolas Brouilly, IBDM, AMU-Marseille), member of the France-BioImaging national research infrastructure (ANR-10-INBS-04) and member of the Marseille Imaging Institute, an Excellence Initiative of Aix Marseille University A*MIDEX, a French “Investissements d’Avenir” programme (AMX 19 IET 002).

## Supplementary Figures and legends

**Fig. S1.**
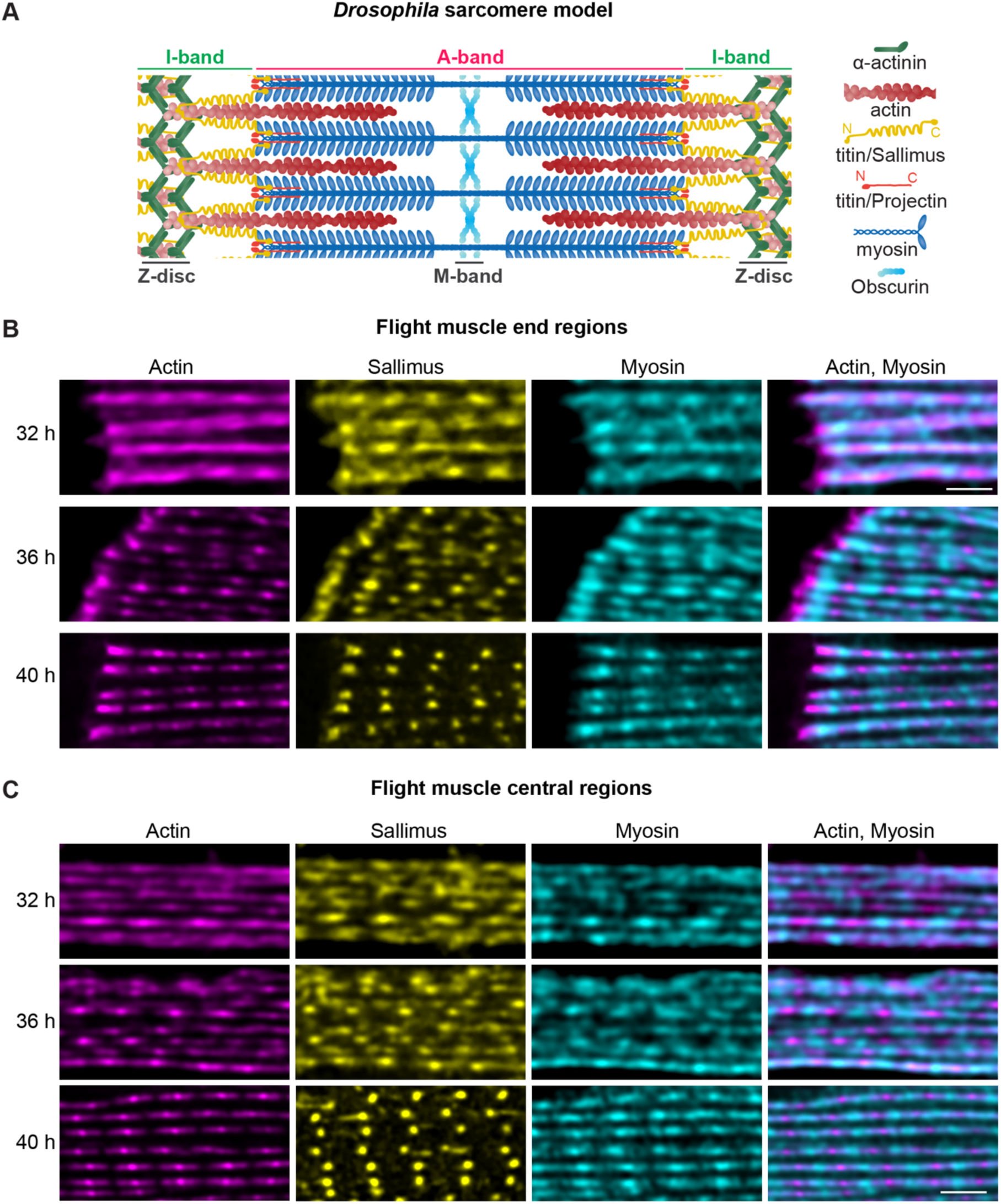
Terminal ends vs. centre of growing myofibrils. **(A)** Schematic of a mature *Drosophila* flight muscle sarcomere bordered by two Z-discs containing α-actinin, which crosslinks polar actin filaments (red). Central bipolar myosin filaments (blue) are cross-linked by Obscurin at the M-band and form the A-band region of the sarcomere. The myosin-free zone is the I-band region. The titin homolog Sallimus (Sls) links the Z-disc to the myosin filaments, the second titin homolog Projectin is located at both ends of the myosin filaments. Scheme was adapted from(*2*) and includes data from(*19*). **(B, C).** High-resolution confocal images of flight muscle myofibrils at 32 h, 36 h and 40 h APF stained for actin (phalloidin in red), Sallimus (Sls-Nano2 in yellow), myosin (anti-Mhc in cyan) focusing on the myofibrils ends in (B) or central regions in (C). Scale bars are 2 µm. Note the periodic patterns of the sarcomeric components, which become more regular over time. Note the slightly higher amounts of actin and Sls at the terminal Z-discs (B). No irregular sarcomeres are present at the terminal ends.

**Fig. S2.**
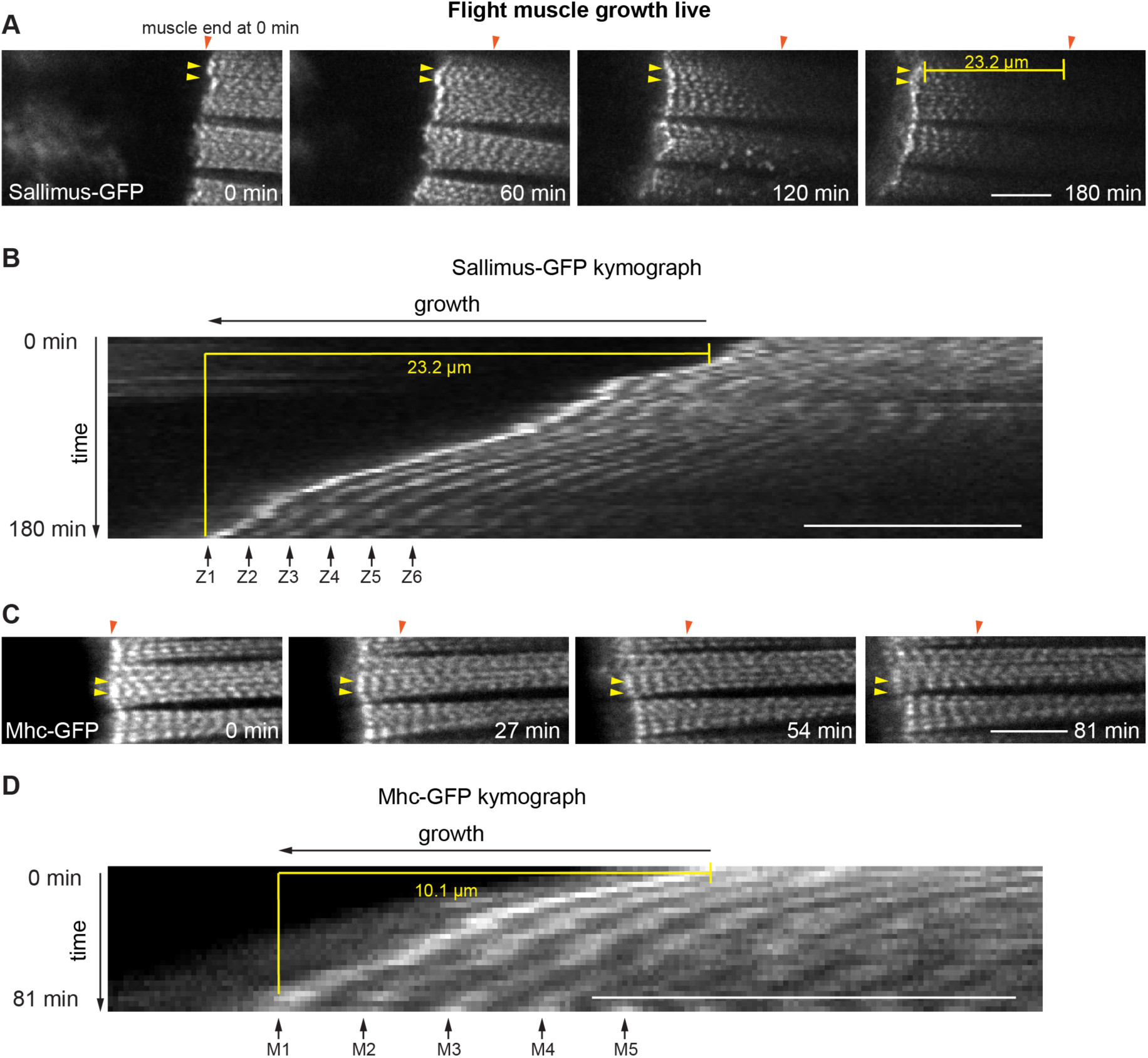
Flight muscle end growth live. **(A, B)** About 34 h APF pupa expressing Sls-GFP with a focus on the anterior flight muscle end. Stills from Movie S1 are shown (A), and kymograph of a selected region is shown (B). Note that the myofibril ends, marked with yellow arrowheads, move to the left while muscle fibre length increases and thus move away from the reference point marked with the orange arrowhead. No enrichment of sarcomere addition at the ends is seen. **(C, D)** About 36 h APF pupa expressing Mhc-GFP with a focus on the anterior flight muscle end. Stills from Movie S2 are shown (C), and kymograph of a selected region is shown (D). Note that the myofibril ends, marked with yellow arrowheads, move to the left while muscle fibre length increases and thus move away from the reference point marked with the orange arrowhead. No enrichment of sarcomere addition is seen. Scale bars are 10 µm.

**Fig. S3.**
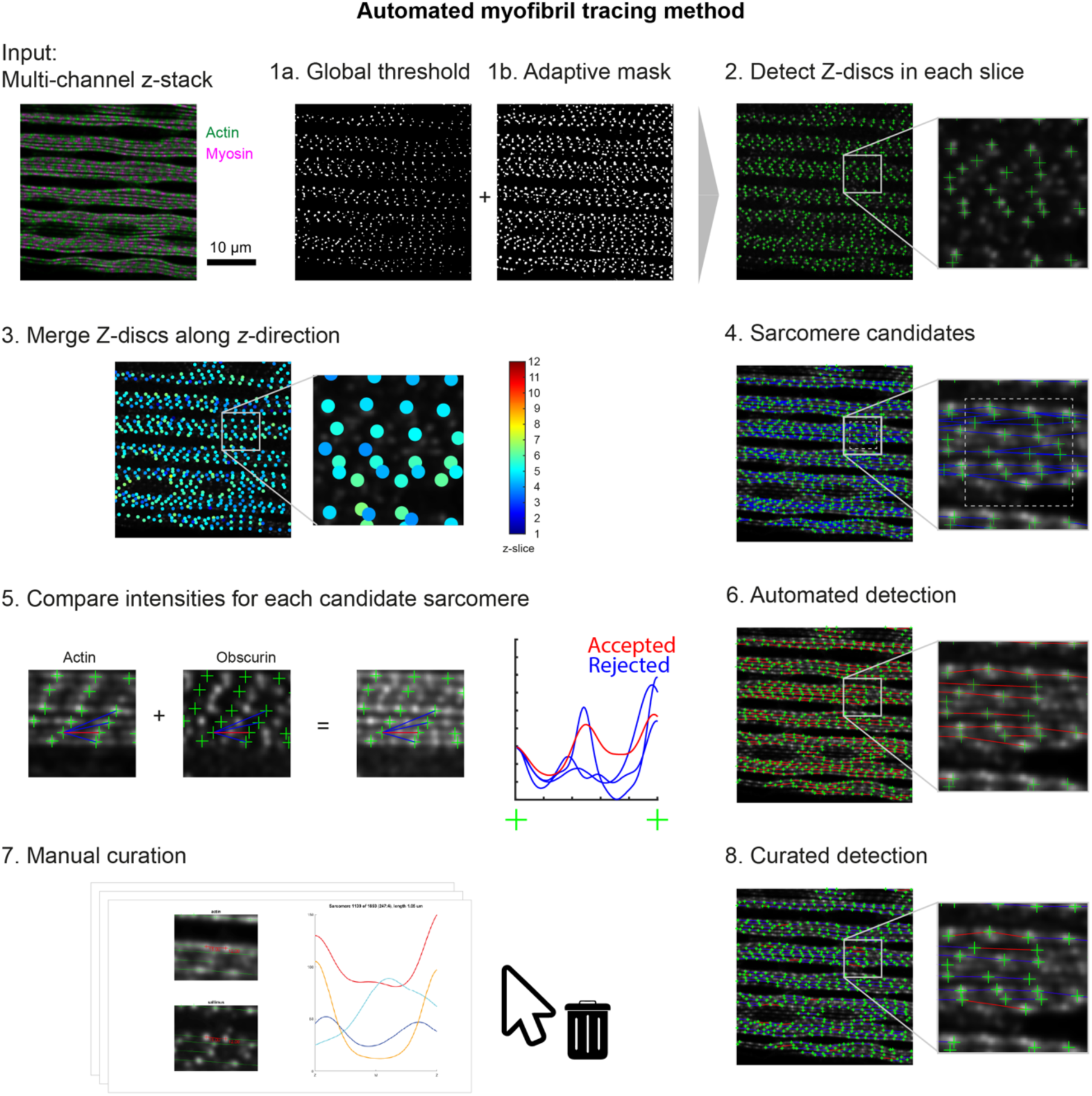
Automated sarcomere detection. **Input:** multi-channel confocal microscopy z-stacks of flight muscles stained for Obscurin, actin, myosin and Sallimus, here represented as typical slice with merge of actin (green) and myosin channels (magenta). **Step 1.** Computation of binary masks from the Sallimus channel using global thresholding (1a) and adaptive thresholding (1b). **Step 2.** Z-disc identification in each z-slice as intensity-weighted centres-of-mass of each connected component of a combined mask (green crosses), obtained as logical AND of the masks from step 1. **Step 3.** For each identified Z-disc, its z-position is determined with sub-voxel resolution by fitting a cubic spline to the Sallimus signal along the z-direction. Identified Z-discs from different slices were merged if their Euclidian distance was smaller than 0.4 µm. Shown are detected Z-discs near selected slice with z-position indicated by colour code. **Step 4.** Identifying sarcomere candidates (blue lines) by connecting pairs of neighbouring Z discs (green crosses) along the myofiber axis within a given tolerance and a maximal length of 3.5 µm. **Step 5.** Automatic selection of true sarcomeres from these candidates, based on the combined fluorescence intensity of actin and Obscurin channels along the sarcomere length. **Step 6.** Result of the automated sarcomere detection. **Step 7.** Manual curation of automated sarcomere detection using a custom-made manual curation tool. **Step 8.** Curated myofibril and sarcomere detection results (red: discarded sarcomeres, blue: retained sarcomeres).

**Fig. S4.**
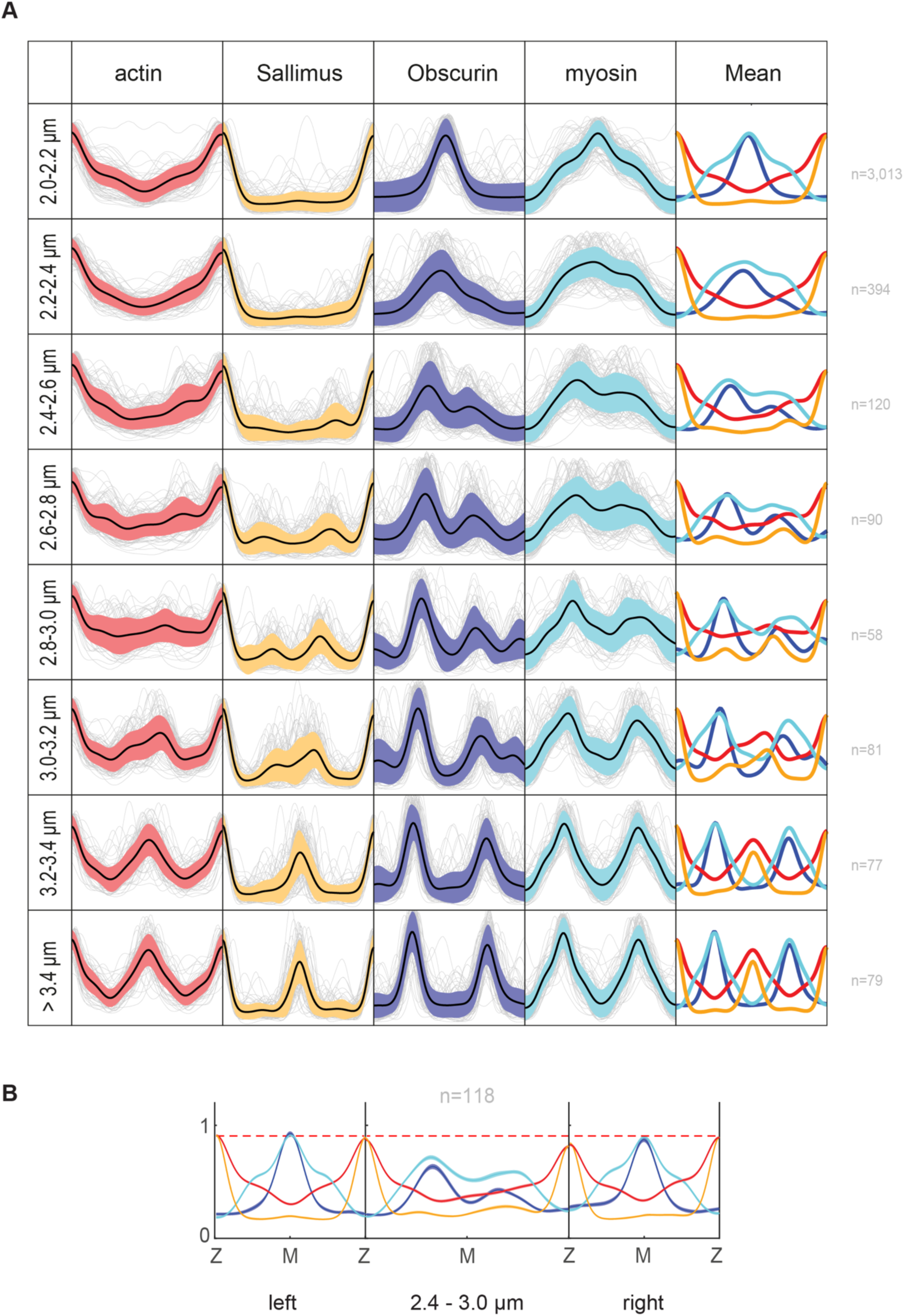
Individual and mean intensity profiles of regular and dividing sarcomeres. **(A)** Grey curves show normalised intensity profiles of individual sarcomeres detected at 40 h APF for actin, Obscurin, Sallimus and myosin, sorted according to sarcomere length (same samples as used for Fig. 2). Black curves and shaded regions depict mean ± s.d.; the right-most column shows the profiles as mean ± s.e.m. **(B)** Intensity profiles for actin, Obscurin, Sallimus and myosin from all 40 h sarcomeres that have a length between 2.4 – 3.0 µm and contain right and left neighbours detected by the algorithm. Note that these neighbours are regular, with a slight reduction in actin and Sls intensities at the shared Z-discs.

**Fig. S5.**
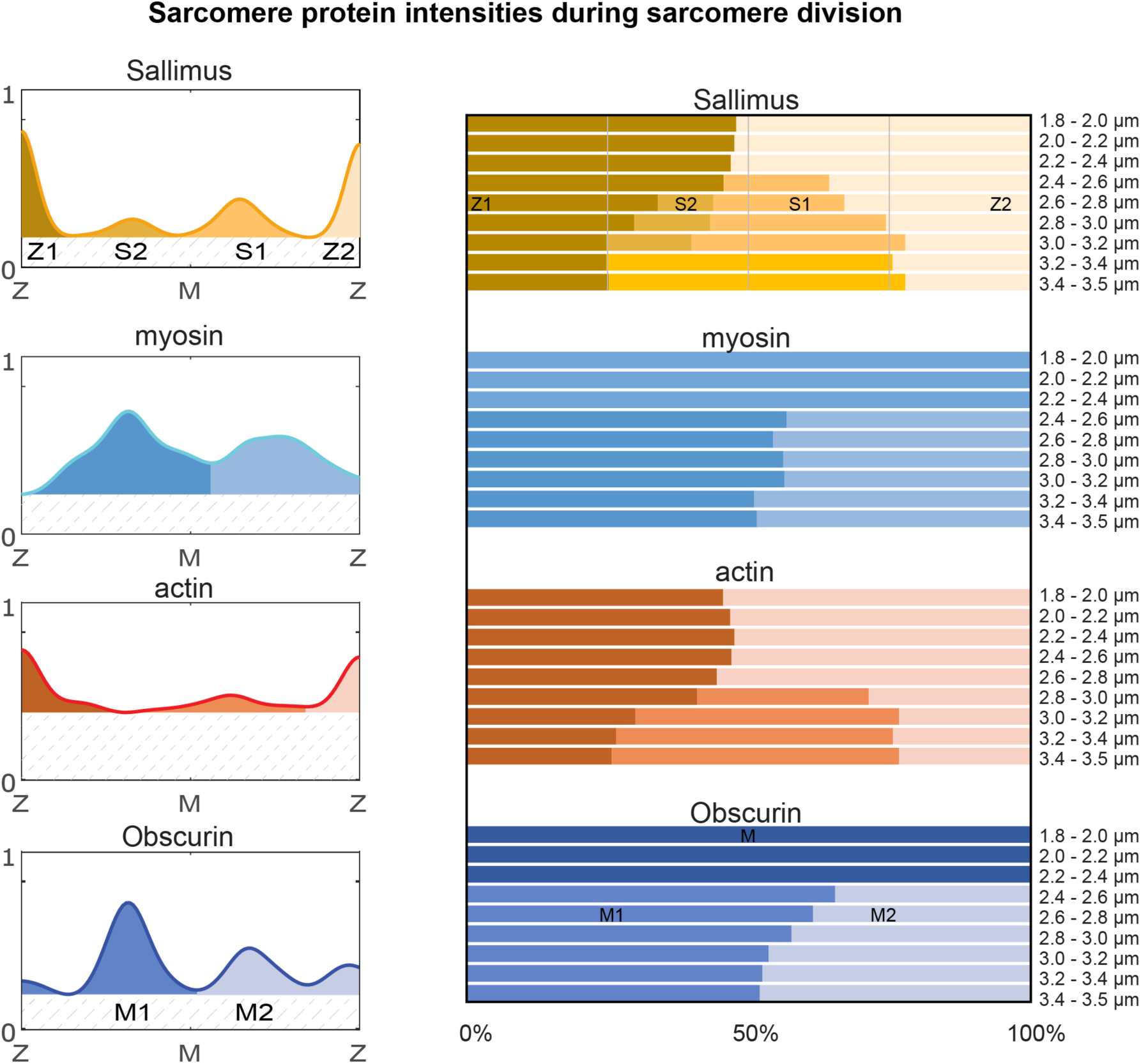
Sarcomere protein intensities as peak areas of mean profiles. Relative areas under the peaks from the mean intensity profiles of sarcomeres within the indicated length bins for Sallimus, myosin, actin, and Obscurin at 40 h APF. Mean profiles were obtained analogously to those shown in Fig. 2D, but using bins as indicated. Different peaks are represented by colours as shown in the example mean profiles to the left. To compute the areas under the peaks, a background intensity given by the minimum of the respective mean profile was subtracted. The relative protein amounts found in each of the peaks for each sarcomere bin size are shown on the right.

**Fig. S6.**
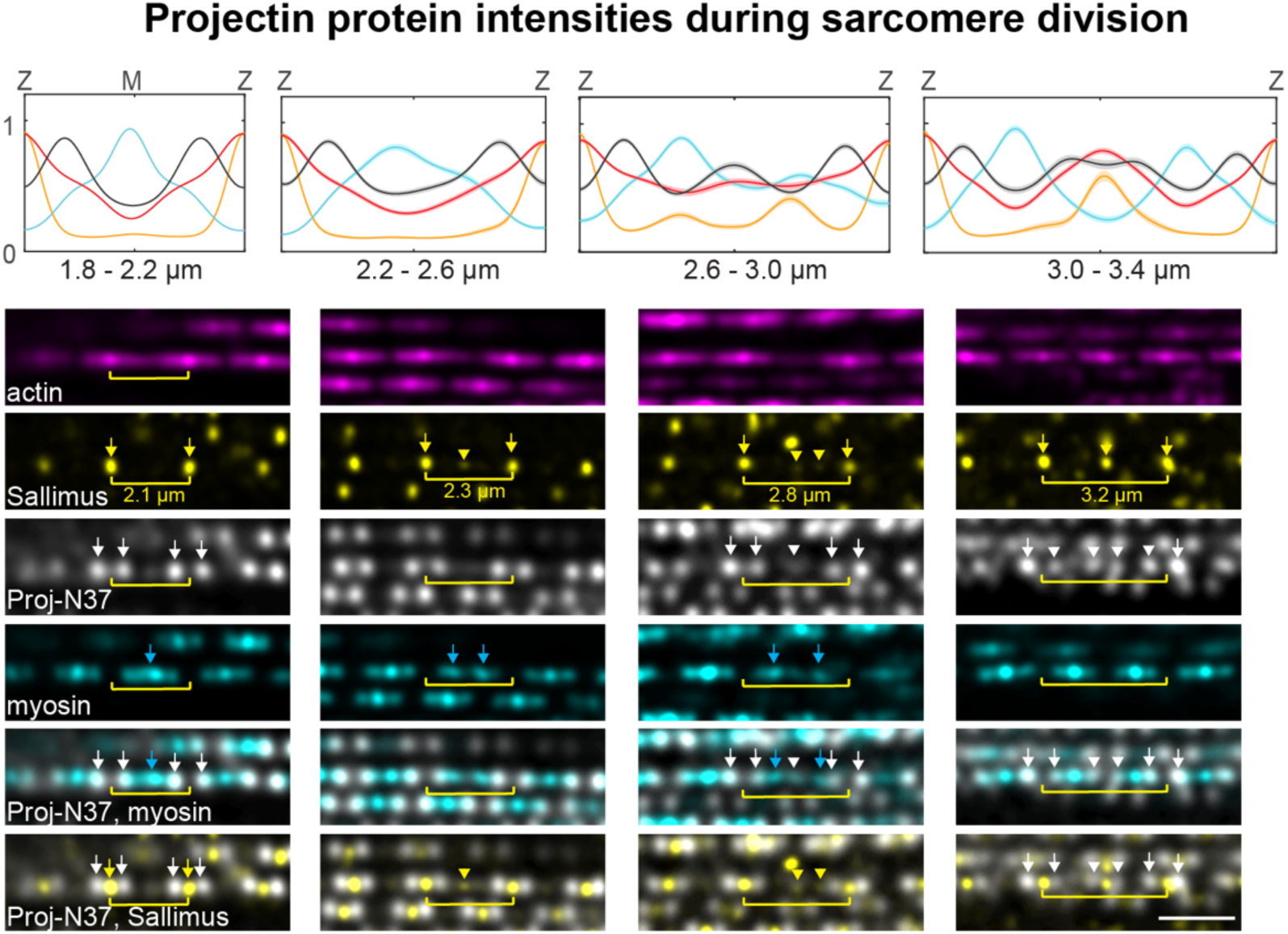
Projectin segregates with myosin during sarcomere division. **Top:** mean intensity profiles of sarcomere proteins within the indicated sarcomere length bins from flight muscles at 40 h APF stained for actin (phalloidin in red), and C-terminus of Projectin (Proj-Nano37 in white), Sallimus (Sls-Nano2 in yellow) and myosin (Mhc-GFP in cyan); mean ± s.e.m. (shaded regions); 5 samples, 1.8 - 2.2 µm: *n* = 5,298 sarcomeres, 2.2 - 2.6 µm: *n* = 35, 2.6 - 3.0 µm: *n* = 55, 3.0 - 3.4 µm: *n* = 41). Below are representative confocal images with highlighted sarcomeres matching the length categories. Note that the two distinct Projectin peaks (white arrows) labelling the ends of the myosin filaments (blue arrows) in regular sarcomeres (1.8 - 2.2 µm) move with the dividing myosin filaments to form two additional new peaks (white arrowheads) next to the newly emerging Z-disc (yellow arrowhead). Scale bar is 2 µm.

**Fig. S7.**
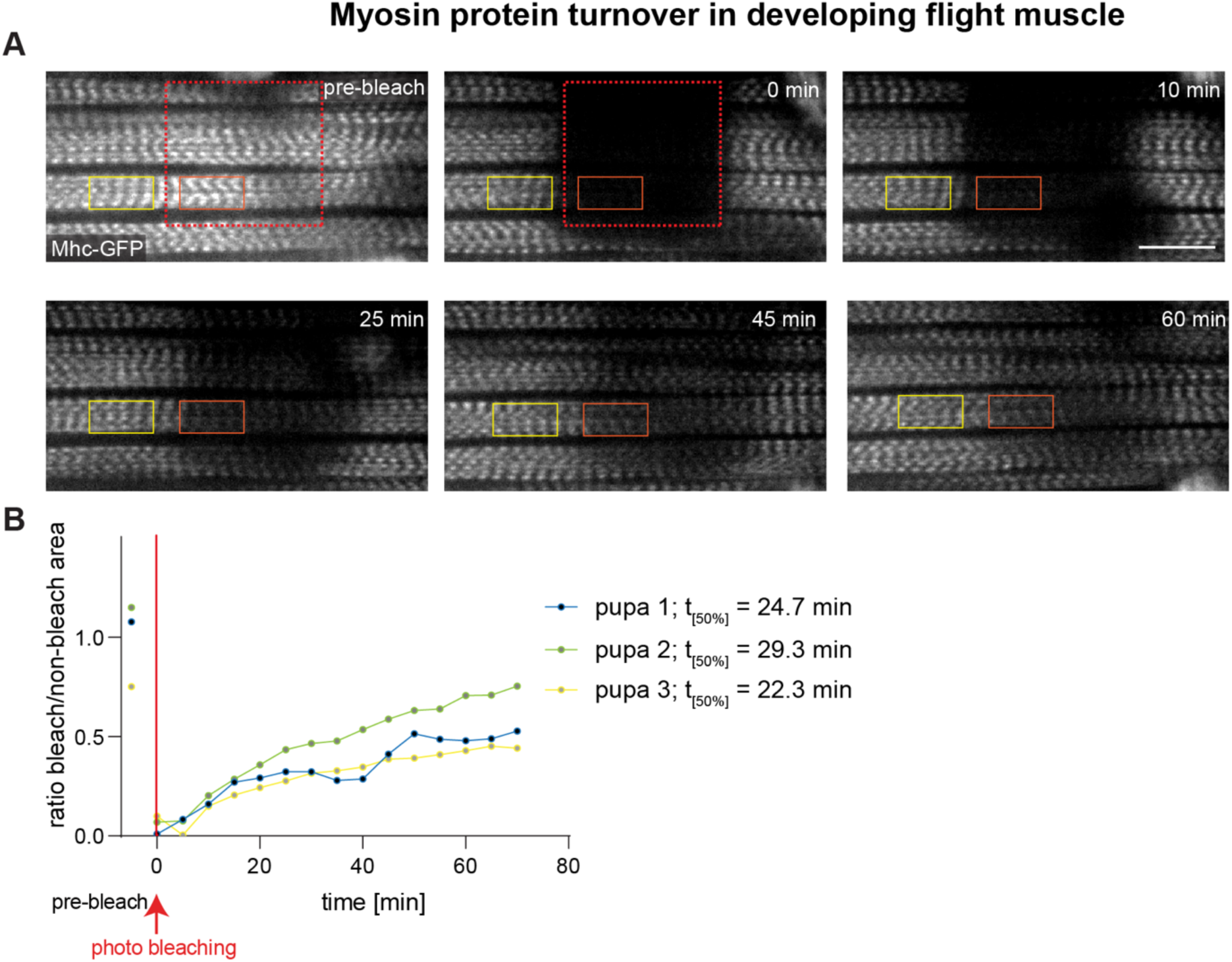
Myosin turnover in flight muscles. **(A)** Stills from 36 h APF flight muscles expressing MhcGFP imaged every 5 min. At 0 min, a square was bleached using the 488 nm laser and recovery of the signal was imaged. Scale bar is 10 µm. (B) Quantification of the Mhc-GFP recovery from 3 representative pupae. T_50%_ recovery time was calculated (see Methods).

**Fig. S8:**
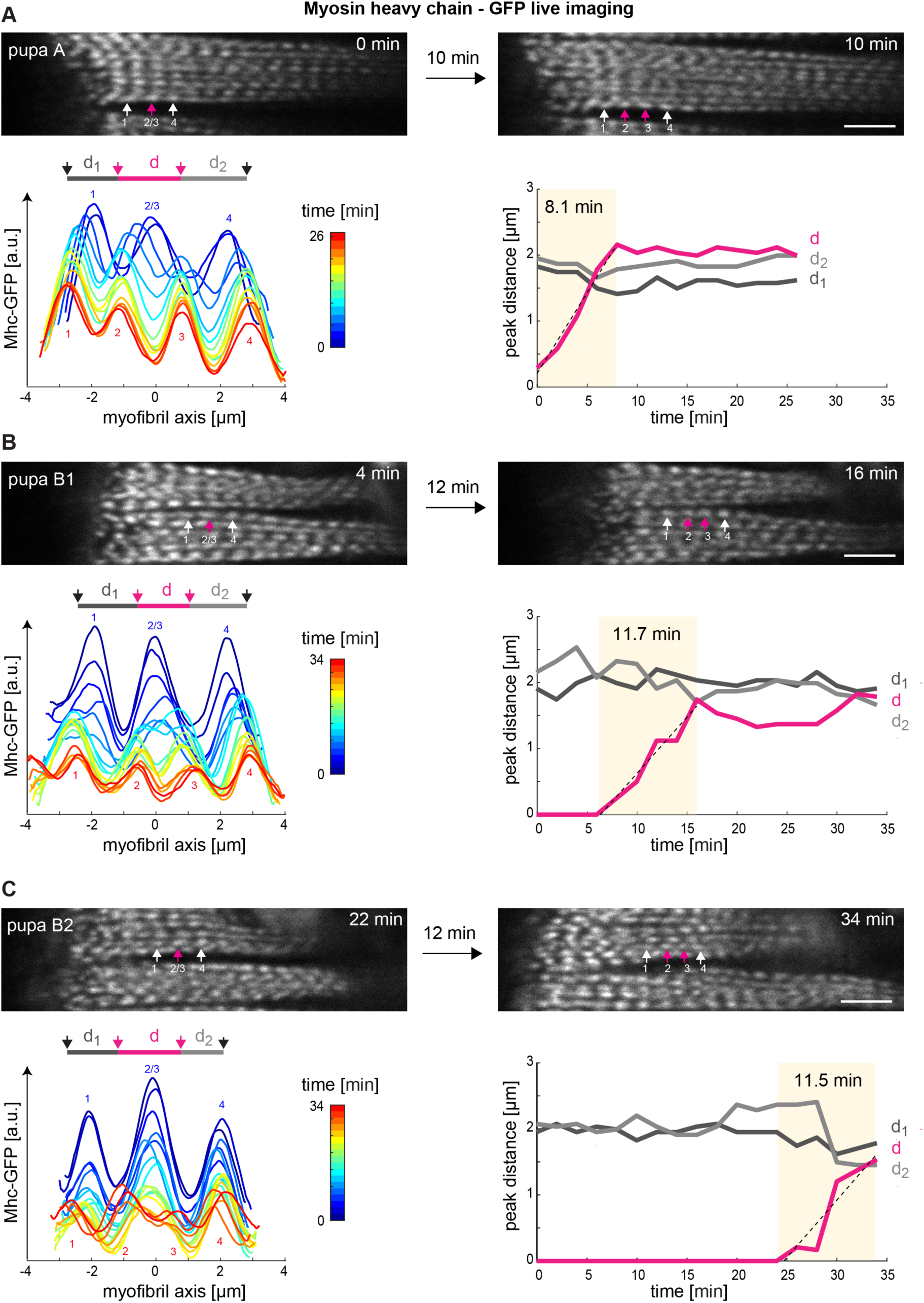
Additional sarcomere division. **(A-C)** Stills from time-lapse movies of Mhc-GFP expressing flight muscles at 36 h APF pupae analogous to Fig. 3 (see Movie S8). In each movie, three sarcomeres labelled 1, 2/3, 4 were manually selected from which the middle one labelled 2/3 will divide into 2 daughters in each movie. Mhc-GFP intensity profiles along the chosen myofibrils are shown below for different colour-coded time points (*left*), together with the distances between the highlighted Mhc-GFP peaks (*right*). Note that the divisions take about 10 to 15 minutes. Scale bars are 5 µm.

**Fig. S9:**
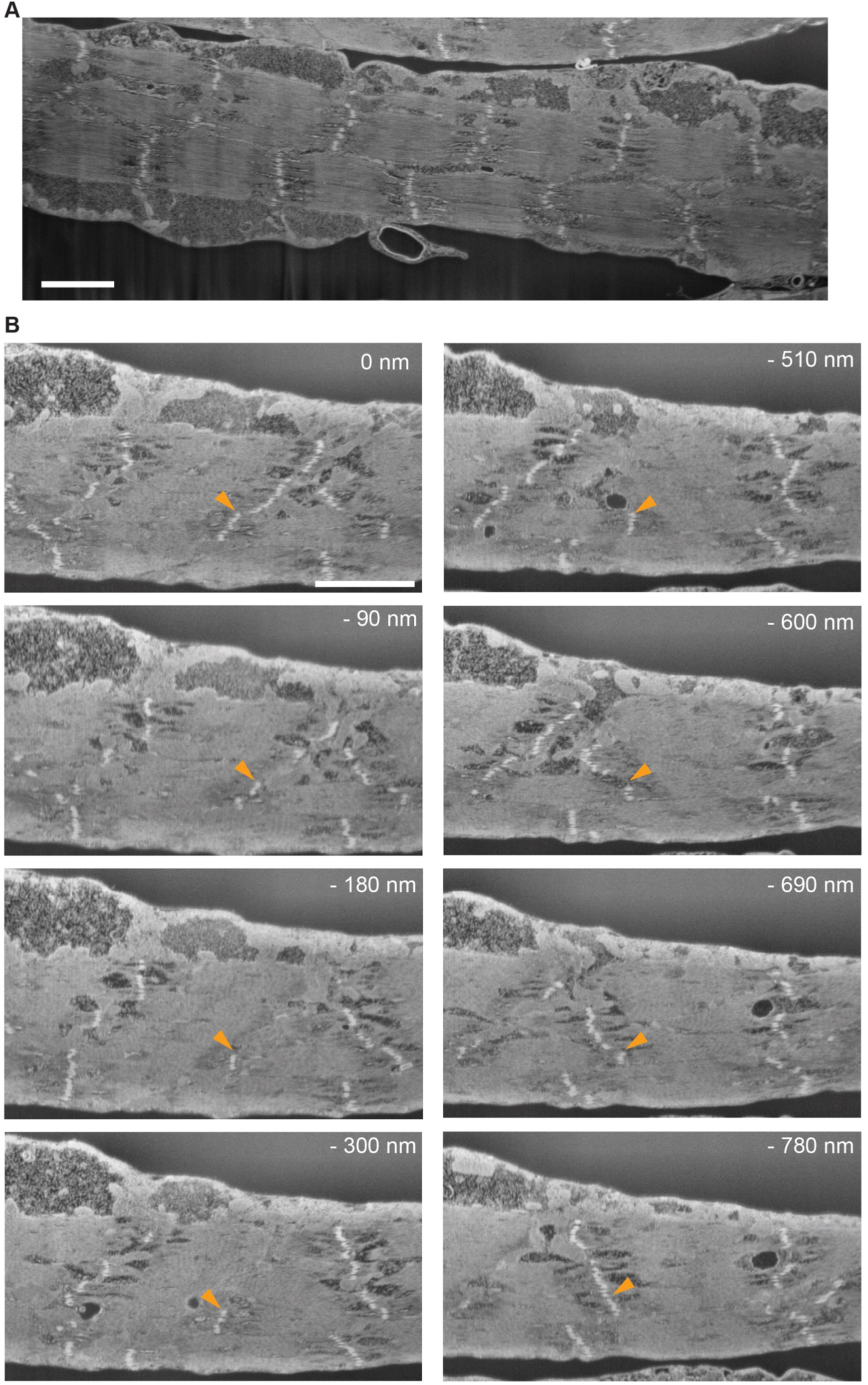
Examples of SEM slices of larval muscles. **(A)** Representative SEM image of a *Drosophila* larval muscle with stacked myofibrils showing organized sarcomeres. (**B**) SEM images from various slices acquired from the same volume, identifying where an isolated Z-disc marked by an orange arrowhead is connected to the Z-disc from an adjacent sarcomere. The relative position of each slice in the volume is indicated relative to the top slice. Scale bars are 5 µm.

**Fig. S10:**
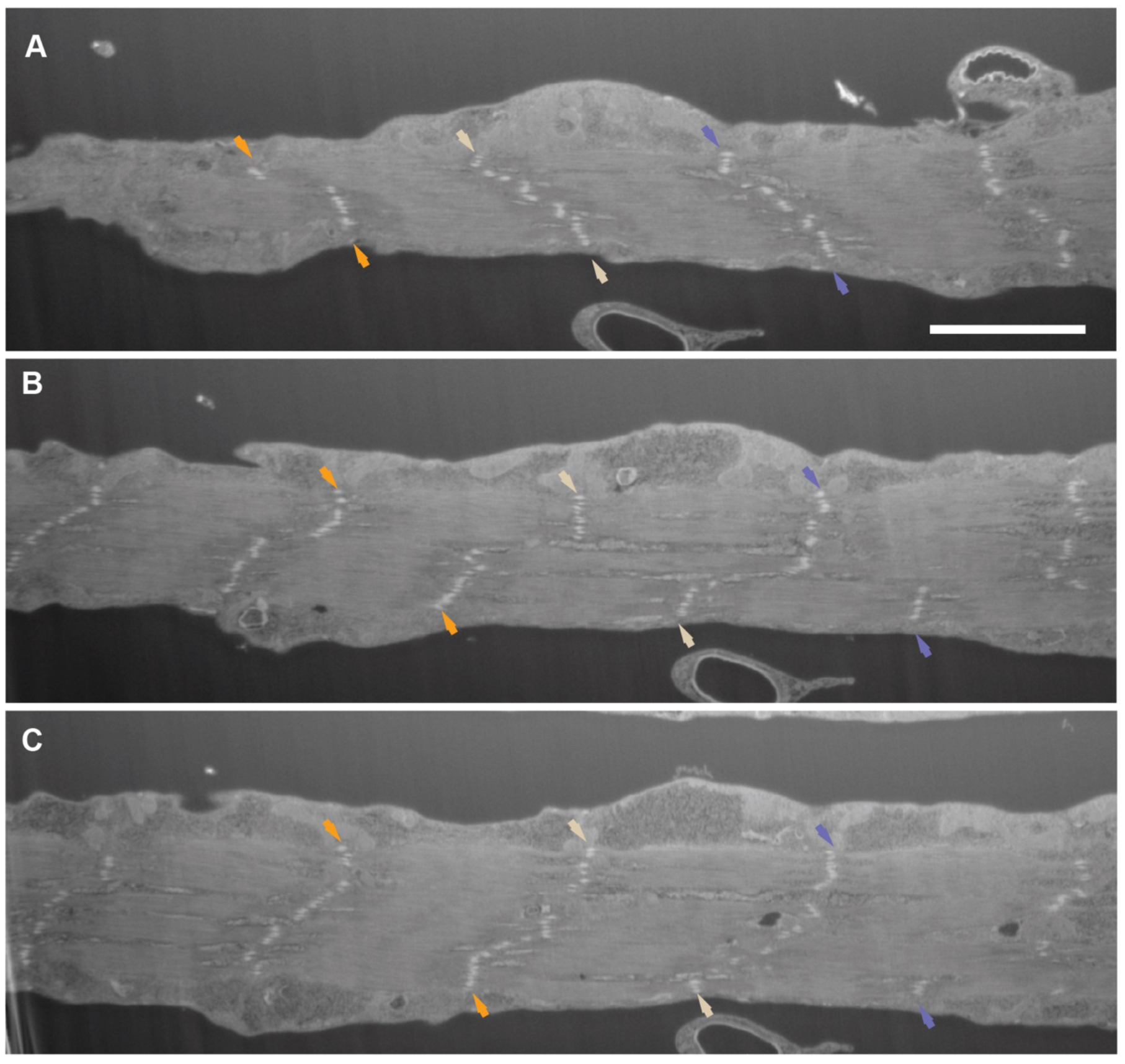
SEM reconstruction of larval muscles. **(A)** SEM image at the surface of a larval muscle. Coloured arrowheads indicate ends of Z-discs that are in the same register at the muscle edge. (**B**) SEM image from 300 nm deeper in the muscle, at which Z-discs have branched in a different direction. (**C**) SEM image from a further 240 nm deeper in the muscle, at which Z-disc connections are reformed between adjacent sarcomeres. This volume was reconstructed in Fig. 4A and Movie S9. Scale bar is 7 µm.

**Fig. S11:**
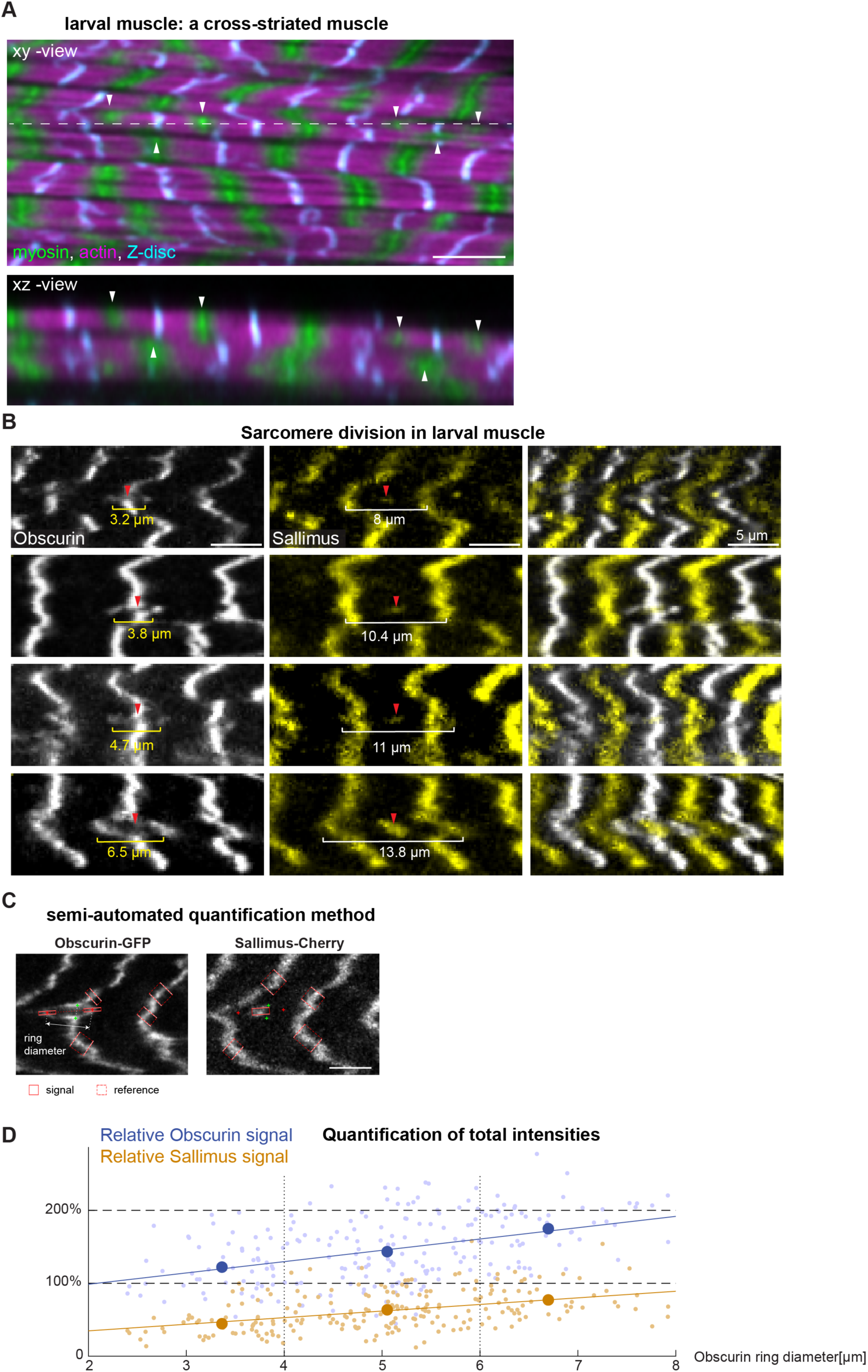
Sarcomeres in living larval muscles. **(A**) High-resolution image of a fixed larval muscle expressing Mhc-GFP (green) stained for actin (phalloidin, magenta) and Sls (Sls-Nano2 cyan). xy-plane shown at the top and reconstructed xz-view at the bottom. Note that myosin stacks (green, marked by white arrowheads) can be separated in xy but remain connected in z. See also Movie S10. (**B**) Living larvae expressing Obscurin-GFP (white) and Sallimus-mCherry (yellow) in muscles. From top to bottom, dividing sarcomeres of increasing ring sizes were sorted. Note that the cross-striated myosin filament stacks segregate, while new Z-disc material (Sls) continues to be incorporated at increasing sarcomere length. **(C)** Illustration of the semi-automated quantification method used for (D). Shown is a single slice of a two-channel z-stack with Obscurin-GFP (left) and Sallimus-mCherry (right); green crosses: manually determined junction points joining the Obscurin ring to M-bands above or below; red crosses: marking the outermost points of the ring; solids red rectangles: regions for line scans of Obscurin and Sls signal; dashed red rectangles: regions for reference line scans used for normalisation. (**D**) Intensity quantifications of dividing sarcomeres sorted by Obscurin-GFP ring width. Note that the relative intensity of Obscurin doubles during the division of one A-band to two A-bands and the amount of Sls increases to reach the amount of a regular sarcomere (gold: Sls, blue: Obscurin; small symbols: individual ‘rings’, large symbols: binned data with bin boundaries indicated by dashed lines; solid lines: linear regression; F-test versus constant model: Obscurin: *p*=1.3 10^-11^, Sls: *p*=8.0 10^-12^). All scale bars are 5 µm.

**Fig. S12:**
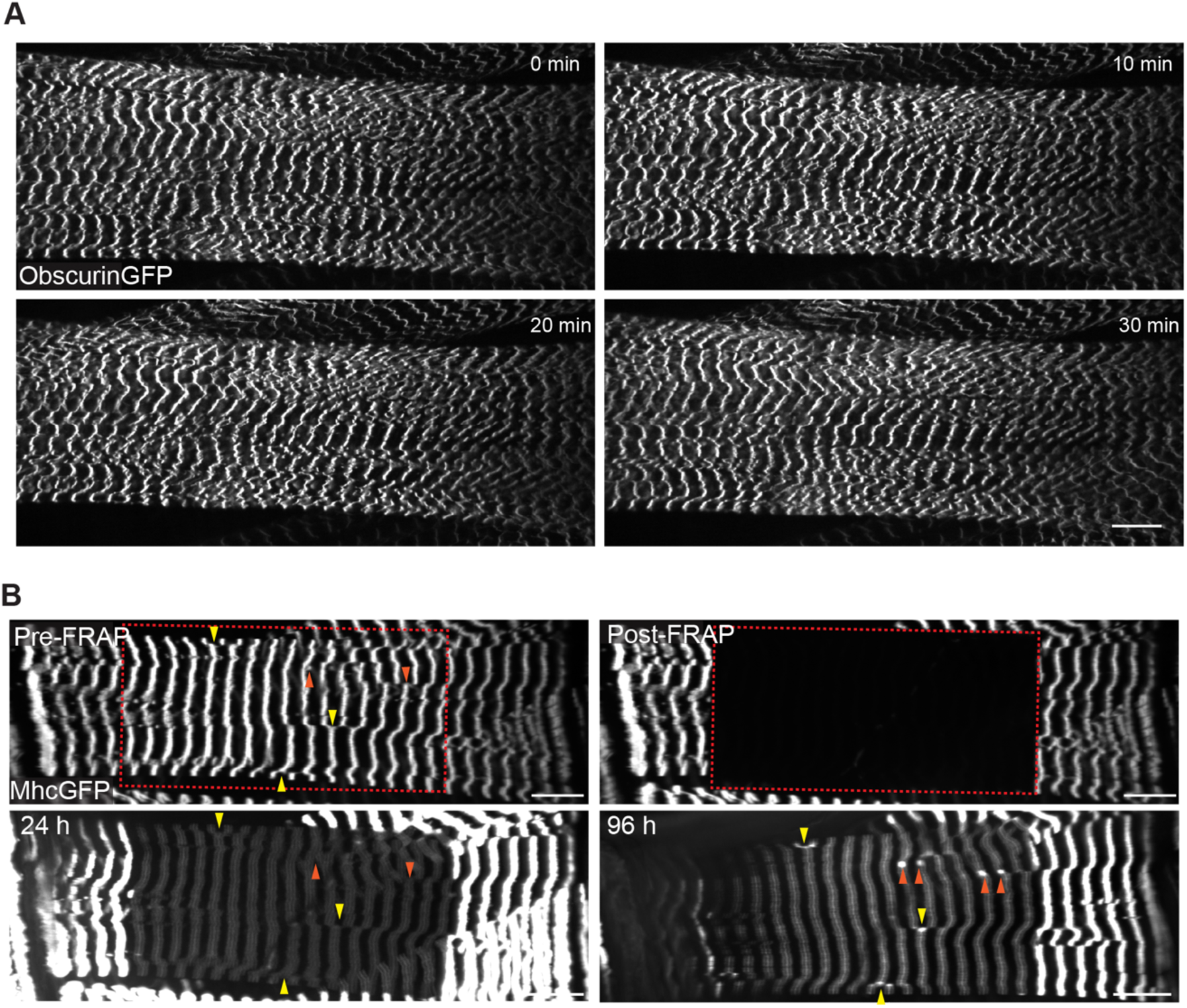
Live imaging of larval muscle and FRAP. **(A)** Stills from a 30-min movie of a living anaesthetised L3 larva expressing Obscurin-GFP to label M-bands. Note that M-bands show little dynamics and sarcomere number remains constant (see Movie S11). (**B**) Mhc-GFP in living L3 larval muscle (dorsal oblique 2, DO2), before FRAP (left, red square) and after FRAP (right), as well as after 24 h (left, bottom) and 96 h (right bottom) of recovery in food. Note the incorporation of new Mhc-GFP in pairs marked by orange arrowheads or at myosin ‘zippers’ (yellow arrowheads). Scale bars are 20 µm.

---

**Movie S1. Live imaging of Sls-GFP at flight muscle ends.** Live imaging of 34 h APF pupa expressing Sls-GFP with a focus on the anterior flight muscle end, imaged every 3 minutes. Arrowheads follow the first 3 sarcomeres over time.

**Movie S2. Live imaging of Mhc-GFP at flight muscle ends.** Live imaging of 36 h APF pupa expressing Mhc-GFP with a focus on the anterior flight muscle end, imaged every 3 minutes. Arrowheads follow the first 3 sarcomeres over time.

**Movie S3. Three-dimensional visualisation of traced myofibrils at 40 h APF.** Animation of traced myofibrils identified in a *z*-stack of 40 h APF flight muscles stained with Sallimus in green (Sls-Nano2), myosin in blue (Mhc-GFP) and actin in red (phalloidin). Grey lines indicate the detected sarcomeres between 2 green dots.

**Movie S4. Three-dimensional visualisation of traced myofibrils at 48 h APF.** Similar to Movie S3 using a 48 h APF flight muscle sample.

**Movie S5: Animation of sarcomere division.** Animation of a dividing sarcomere using the model shown in Fig. 2E. Note that the daughter 1 sarcomere (Z1, M1, S1) segregates from daughter 2 sarcomere (S2, M2, Z2). At the end, S1 and S2 fuse and establish the new Z-disc.

**Movie S6. Live imaging of Mhc-GFP turnover in flight muscles.** Live imaging of Mhc-GFP expressing flight muscles at 36 h APF with spinning disc microscopy, imaged every 5 min. The white rectangle area was bleached and Mhc-GFP recovery was followed.

**Movie S7. Imaging and tracking of A-band division live.** Two-photon microscopy live imaging of Mhc-GFP expressing flight muscles at 36 h APF. Myosin stacks were automatically tracked along one myofibril marked with black crosses. The one marked with a red cross divides into two myosin stacks, starting at 0 minutes. These data were used to quantify distances in Fig. 3.

**Movie S8. Imaging of A-band divisions live.** Two-photon microscopy live imaging of two different pupae expressing flight muscles at 36 h APF. Dividing myosin stacks and their immediate neighbours were manually marked with arrowheads and followed over time. Note myosin filament stack divisions occur anywhere in the muscle cell.

**Movie S9. Three-dimensional reconstruction of larval muscle with SEM.** Scanning electron microscopy segmentation of larval Z-discs in a 3D volume starting from the muscle surface using the slices shown in Fig. S10.

**Movie S10. Three-dimensional view of larval muscle.** Z-stack of high-resolution images of a fixed larval muscle expressing Mhc-GFP (green) stained for actin (phalloidin, magenta) and Sls (Sls-Nano2 cyan). Note that M-bands and Z-stacks are often segregated in 2D, but stay largely connected in 3D.

**Movie S11. Live imaging of anaesthetised larvae.** Live imaging of an Obscurin-GFP expressing dorsal oblique 2 muscle (DO2) in an anaesthetised larva, imaged every 2 minutes with a spinning disc confocal microscope. Note that M-bands are stable over the entire movie.

**Movie S12. Dynamic recovery of Mhc-GFP after FRAP.** Z-stack of living larva that expresses Mhc-GFP in green and Sls-Cherry in magenta, 7 hours after FRAP recovery with feeding. Note that Mhc-GFP does recover in pairs marked by arrowheads in the individual planes.

**Data S1** – Data related for Fig. 1

**Data S2** – Data related for Fig. 2

**Data S3** – Data related for Fig. 3

**Data S4** – Data related for Fig. 4

## References

1. M. E. Llewellyn, R. P. J. Barretto, S. L. Delp, M. J. Schnitzer, Minimally invasive high-speed imaging of sarcomere contractile dynamics in mice and humans. Nature 454, 784–788 (2008).

2. N. M. Luis, F. Schnorrer, Mechanobiology of muscle and myofibril morphogenesis. Cells Dev. 168, 203760 (2021).

3. W. A. Linke, Titin Gene and Protein Functions in Passive and Active Muscle. Annu. Rev. Physiol. 80, 389–411 (2018).

4. T. R. Heallen, Z. A. Kadow, J. Wang, J. F. Martin, Determinants of Cardiac Growth and Size. Cold Spring Harb. Perspect. Biol. 12, a037150 (2020).

5. G. Kardon, Development of the musculoskeletal system: meeting the neighbors. Development 138, 2855–2859 (2011).

6. Q. Mao, A. Acharya, A. Rodríguez-delaRosa, F. Marchiano, B. Dehapiot, Z. Al Tanoury, J. Rao, M. Díaz-Cuadros, A. Mansur, E. Wagner, C. Chardes, V. Gupta, P.-F. Lenne, B. H. Habermann, O. Theodoly, O. Pourquié, F. Schnorrer, Tension-driven multi-scale self-organisation in human iPSC-derived muscle fibers. eLife 11, e76649 (2022).

7. D. J. Dix, B. R. Eisenberg, Myosin mRNA accumulation and myofibrillogenesis at the myotendinous junction of stretched muscle fibers. J. Cell Biol. 111, 1885–1894 (1990).

8. P. E. Williams, G. Goldspink, The effect of immobilization on the longitudinal growth of striated muscle fibres. J. Anat. 116, 45–55 (1973).

9. H. J. Green, A. G. Griffiths, J. Ylänne, N. H. Brown, Novel functions for integrin-associated proteins revealed by analysis of myofibril attachment in Drosophila. eLife 7, e35783 (2018).

10. K. M. Vakaloglou, G. Chrysanthis, M. A. Rapsomaniki, Z. Lygerou, C. G. Zervas, IPP Complex Reinforces Adhesion by Relaying Tension-Dependent Signals to Inhibit Integrin Turnover. Cell Rep. 14, 2668–2682 (2016).

11. M. Balakrishnan, S. F. Yu, S. M. Chin, D. B. Soffar, S. E. Windner, B. L. Goode, M. K. Baylies, Cofilin Loss in Drosophila Muscles Contributes to Muscle Weakness through Defective Sarcomerogenesis during Muscle Growth. Cell Rep. 32, 107893 (2020).

12. M. Ganassi, S. Badodi, K. Wanders, P. S. Zammit, S. M. Hughes, Myogenin is an essential regulator of adult myofibre growth and muscle stem cell homeostasis. eLife 9, e60445 (2020).

13. S. B. Lemke, F. Schnorrer, Mechanical forces during muscle development. Mech. Dev. 144, 92–101 (2017).

14. M. L. Spletter, C. Barz, A. Yeroslaviz, X. Zhang, S. B. Lemke, A. Bonnard, E. Brunner, G. Cardone, K. Basler, B. H. Habermann, F. Schnorrer, A transcriptomics resource reveals a transcriptional transition during ordered sarcomere morphogenesis in flight muscle. eLife 7, e34058 (2018).

15. M. Weitkunat, A. Kaya-Çopur, S. W. Grill, F. Schnorrer, Tension and Force-Resistant Attachment Are Essential for Myofibrillogenesis in Drosophila Flight Muscle. Curr. Biol. 24, 705–716 (2014).

16. A. Kaya-Çopur, F. Marchiano, M. Y. Hein, D. Alpern, J. Russeil, N. M. Luis, M. Mann, B. Deplancke, B. H. Habermann, F. Schnorrer, The Hippo pathway controls myofibril assembly and muscle fiber growth by regulating sarcomeric gene expression. eLife 10, e63726 (2021).

17. Z. Orfanos, K. Leonard, C. Elliott, A. Katzemich, B. Bullard, J. Sparrow, Sallimus and the Dynamics of Sarcomere Assembly in Drosophila Flight Muscles. J. Mol. Biol. 427, 2151–2158 (2015).

18. O. Loison, M. Weitkunat, A. Kaya-Çopur, C. Nascimento Alves, T. Matzat, M. L. Spletter, S. Luschnig, S. Brasselet, P.-F. Lenne, F. Schnorrer, Polarization-resolved microscopy reveals a muscle myosin motor-independent mechanism of molecular actin ordering during sarcomere maturation. PLOS Biol. 16, e2004718 (2018).

19. F. Schueder, P. Mangeol, E. H. Chan, R. Rees, J. Schünemann, R. Jungmann, D. Görlich, F. Schnorrer, Nanobodies combined with DNA-PAINT super-resolution reveal a staggered titin nanoarchitecture in flight muscles. eLife 12, e79344 (2023).

20. Z. Orfanos, J. C. Sparrow, Myosin isoform switching during assembly of the *Drosophila* flight muscle thick filament lattice. J. Cell Sci. 126, 139–148 (2013).

21. C. Schönbauer, J. Distler, N. Jährling, M. Radolf, H.-U. Dodt, M. Frasch, F. Schnorrer, Spalt mediates an evolutionarily conserved switch to fibrillar muscle fate in insects. Nature 479, 406–409 (2011).

22. F. Demontis, N. Perrimon, Integration of Insulin receptor/Foxo signaling and dMyc activity during muscle growth regulates body size in *Drosophila*. Development 136, 983–993 (2009).

23. V. Loreau, R. Rees, E. H. Chan, W. Taxer, K. Gregor, B. Mußil, C. Pitaval, N. M. Luis, P. Mangeol, F. Schnorrer, D. Görlich, A nanobody toolbox to investigate localisation and dynamics of Drosophila titins and other key sarcomeric proteins. eLife 12, e79343 (2023).

24. Y. Li, A. L. Hessel, A. Unger, D. Ing, J. Recker, F. Koser, J. K. Freundt, W. A. Linke, Graded titin cleavage progressively reduces tension and uncovers the source of A-band stability in contracting muscle. eLife 9, e64107 (2020).

25. V. Loreau, W. Koolhaas, E. H. Chan, P. De Bossier, N. Brouilly, S. Avosani, A. Sane, C. Pitaval, S. Reiter, N. M. Luis, P. Mangeol, A. C. Von Philipsborn, J.-F. Rupprecht, D. Goerlich, B. H. Habermann, F. Schnorrer, Titin-dependent biomechanical feedback tailors sarcomeres to specialised muscle functions in insects. [Preprint] (2024). 10.1101/2024.09.30.615857.

26. J. A. Rivas-Pardo, Y. Li, Z. Mártonfalvi, R. Tapia-Rojo, A. Unger, Á. Fernández-Trasancos, E. Herrero-Galán, D. Velázquez-Carreras, J. M. Fernández, W. A. Linke, J. Alegre-Cebollada, A HaloTag-TEV genetic cassette for mechanical phenotyping of proteins from tissues. Nat. Commun. 11, 2060 (2020).

27. S. S. Jahromi, M. P. Charlton, Transverse sarcomere splitting. A possible means of longitudinal growth in crab muscles. J. Cell Biol. 80, 736–742 (1979).

28. M. Gautel, The sarcomeric cytoskeleton: who picks up the strain? Curr. Opin. Cell Biol. 23, 39–46 (2011).

29. P. Young, E. Ehler, M. Gautel, Obscurin, a giant sarcomeric Rho guanine nucleotide exchange factor protein involved in sarcomere assembly. J. Cell Biol. 154, 123–136 (2001).

30. D. Tamborrini, Z. Wang, T. Wagner, S. Tacke, M. Stabrin, M. Grange, A. L. Kho, M. Rees, P. Bennett, M. Gautel, S. Raunser, Structure of the native myosin filament in the relaxed cardiac sarcomere. Nature, 1–9 (2023).

31. L. Carlsson, J.-G. Yu, M. Moza, O. Carpén, L.-E. Thornell, Myotilin – a prominent marker of myofibrillar remodelling. Neuromuscul. Disord. 17, 61–68 (2007).

32. J.-G. Yu, B. Russell, Cardiomyocyte Remodeling and Sarcomere Addition after Uniaxial Static Strain In Vitro. J. Histochem. Cytochem. 53, 839–844 (2005).

33. J.-G. Yu, L.-E. Thornell, Desmin and actin alterations in human muscles affected by delayed onset muscle soreness: a high resolution immunocytochemical study. Histochem. Cell Biol. 118, 171–179 (2002).

34. J.-G. Yu, L. Carlsson, L.-E. Thornell, Evidence for myofibril remodeling as opposed to myofibril damage in human muscles with DOMS: an ultrastructural and immunoelectron microscopic study. Histochem. Cell Biol. 121, 219–227 (2004).

35. R. Lynn, D. L. Morgan, Decline running produces more sarcomeres in rat vastus intermedius muscle fibers than does incline running. J. Appl. Physiol. 77, 1439–1444 (1994).

36. D. L. Morgan, J. A. Talbot, The addition of sarcomeres in series is the main protective mechanism following eccentric exercise. J. Mech. Med. Biol. 02, 421–431 (2002).

37. J. Avellaneda, C. Rodier, F. Daian, N. Brouilly, T. Rival, N. M. Luis, F. Schnorrer, Myofibril and mitochondria morphogenesis are coordinated by a mechanical feedback mechanism in muscle. Nat. Commun. 12, 2091 (2021).

38. M. Sarov, C. Barz, H. Jambor, M. Y. Hein, C. Schmied, D. Suchold, B. Stender, S. Janosch, V. V. Kj, R. Krishnan, A. Krishnamoorthy, I. R. Ferreira, R. K. Ejsmont, K. Finkl, S. Hasse, P. Kämpfer, N. Plewka, E. Vinis, S. Schloissnig, E. Knust, V. Hartenstein, M. Mann, M. Ramaswami, K. VijayRaghavan, P. Tomancak, F. Schnorrer, A genome-wide resource for the analysis of protein localisation in Drosophila. eLife 5, e12068 (2016).

39. M. Weitkunat, F. Schnorrer, A guide to study Drosophila muscle biology. Methods 68, 2– 14 (2014).

40. D. Strumpf, T. Volk, Kakapo, a Novel Cytoskeletal-associated Protein Is Essential for the Restricted Localization of the Neuregulin-like Factor, Vein, at the Muscle–Tendon Junction Site. J. Cell Biol. 143, 1259–1270 (1998).

41. S. B. Lemke, F. Schnorrer, In Vivo Imaging of Muscle-tendon Morphogenesis in Drosophila Pupae. J. Vis. Exp., 57312 (2018).

42. P. Kakanj, S. A. Eming, L. Partridge, M. Leptin, Long-term in vivo imaging of Drosophila larvae. Nat. Protoc. 15, 1158–1187 (2020).

43. T. J. Deerinck, E. A. Bushong, M. H. Ellisman, A. Thor, Preparation of Biological Tissues for Serial Block Face Scanning Electron Microscopy (SBEM). (2022).

44. J. Schindelin, I. Arganda-Carreras, E. Frise, V. Kaynig, M. Longair, T. Pietzsch, S. Preibisch, C. Rueden, S. Saalfeld, B. Schmid, J.-Y. Tinevez, D. J. White, V. Hartenstein, K. Eliceiri, P. Tomancak, A. Cardona, Fiji: an open-source platform for biological-image analysis. Nat. Methods 9, 676–682 (2012).

45. E. C. Meng, T. D. Goddard, E. F. Pettersen, G. S. Couch, Z. J. Pearson, J. H. Morris, T. E. Ferrin, UCSF CHIMERAX: Tools for structure building and analysis. Protein Sci. 32, e4792 (2023).

